# Mapping genomic regulation of kidney disease and traits through high-resolution and interpretable eQTLs

**DOI:** 10.1101/2022.06.01.494352

**Authors:** Seong Kyu Han, Michelle T. McNulty, Christopher J. Benway, Pei Wen, Anya Greenberg, Ana C. Onuchic-Whitford, NEPTUNE, Parker C. Wilson, Benjamin D. Humphreys, Xiaoquan Wen, Zhe Han, Dongwon Lee, Matthew G. Sampson

## Abstract

Expression quantitative trait locus (eQTL) studies illuminate genomic variants that regulate specific genes and contribute to fine-mapped loci discovered via genome-wide association studies (GWAS). Efforts to maximize their accuracy are ongoing. Using 240 glomerular (GLOM) and 311 tubulointerstitial (TUBE) micro-dissected samples from human kidney biopsies, we discovered 5,371 GLOM and 9,787 TUBE eQTLs by incorporating kidney single-nucleus open chromatin data and transcription start site distance as an “integrative prior” for Bayesian statistical fine mapping. The use of an integrative prior resulted in higher resolution eQTLs illustrated by (1) smaller numbers of variants in credible sets with greater confidence, (2) increased enrichment of partitioned heritability for GWAS of two kidney traits, (3) an increased number of variants colocalized with the GWAS loci, and (4) enrichment of computationally predicted functional regulatory variants. A subset of variants and genes were validated experimentally *in vitro* and using a *Drosophila* nephrocyte model. More broadly, this study demonstrates that tissue-specific eQTL maps informed by single-nucleus open chromatin data have enhanced utility for diverse downstream analyses.

## Introduction

The genomic contributors to kidney diseases and traits extend well beyond rare, pathogenic, exonic variants with large effect sizes that typified the initial discoveries in this area. Focused analyses of individual genes have long illuminated common, non-coding variants whose regulatory effects contribute to their proper function (Arking et al., 2006; Tyburczy et al., 2015). More recently, genome-wide association studies (GWASs) have demonstrated that the heritability of diverse kidney traits and diseases are polygenic and primarily non-coding (Kiryluk et al., 2014; Köttgen et al., 2010; Pollak and Friedman, 2020; Teumer et al., 2019; Wuttke et al., 2019; Xie et al., 2020). Thus, whether to deeply understand a single disease-related gene or to fine-map GWAS loci, it is necessary to have as precise an understanding of the genetic control of gene expression as possible. A high-resolution expression quantitative trait loci (eQTL) map of the kidney can contribute greatly to this need.

eQTLs can identify variants associated with gene expression (“eSNPs”) and their target genes (“eGenes”) across individuals in a tissue-, and more recently cell-, informed manner (Kim-Hellmuth et al., 2020; Ongen et al., 2017). In particular, GWAS fine mapping of diseases and traits has been aided by these eQTL maps given that the non-coding nature of most GWAS variants and linkage disequilibrium (LD) preclude their direct interpretation (Cano-Gamez and Trynka, 2020). Beyond eQTLs, annotations from complementary genomic experiments (e.g., open chromatin peaks) can provide further refinement by identifying functional regions within the disease- and trait-associated loci. Previously, prioritizing GWAS SNPs using eQTL and other functional annotations was achieved by simply overlapping these datasets (*post hoc* “lookups”). However, investigators have begun to develop new approaches to empower more precise GWAS fine-mapping, particularly by building more high-resolution eQTL datasets (Cano-Gamez and Trynka, 2020).

One method to improve eQTL mapping is to incorporate single-cell data. In a recent study, investigators predicted kidney cell-type interacting eQTLs by applying *in silico* deconvolution methods using a reference single-cell gene expression dataset and subsequently built a cell fraction-informed eQTL model from bulk tissue (Sheng et al., 2021). This cell-type-informed eQTL data was then co-analyzed with assay for transposase-accessible chromatin using sequencing (ATAC-seq) data *post hoc* to identify overlaps between specific eSNPs and open chromatin.

Another approach to improve the resolution of eQTL maps is to incorporate functionally-informed SNP annotations in the fine-mapping procedure (Gaffney et al., 2012). As a proof of principle, early studies showed the enrichment of regulatory annotations in eQTLs from lymphoblastoid cell lines. Then, they showed that integrating this information into a Bayesian fine-mapping framework could improve the eQTL discovery and fine-mapping resolution (Gaffney et al., 2012; Wen et al., 2015). Similarly, a recent study demonstrated increased statistical fine-mapping accuracy of eQTLs by assigning weights to SNPs using priors derived from diverse functional annotations for the subsequent eQTL analysis (Wang et al., 2021). This approach increases eQTL discovery and loci for downstream consideration that would have been missed using the *post hoc* “lookup” strategy.

In this study, we further extend this approach by incorporating single-nucleus open chromatin data from kidney tissue to fine-map kidney eQTLs. Given that SNPs within open chromatin peaks are more likely to impact transcriptional regulation (Gaffney et al., 2012), we hypothesized that weighting SNPs by this parameter would increase eQTL discovery and fine mapping of putative functional SNPs that otherwise would not be found due to high LD or low allele frequencies. In doing so, we hypothesized that we would gain 1) greater statistical and functional confidence in putative causal eSNPs, 2) additional insight into the regulatory landscape for specific genes related to kidney diseases and traits, and 3) increased discovery and fine-mapping resolution for genome-wide, integrative analyses.

To test this hypothesis, we created a workflow to discover “high-resolution eQTLs” by using single-nucleus open chromatin data from kidney tissue to generate priors for use in a Bayesian multi-SNP eQTL detection and fine-mapping algorithm (**Figure 1**). We applied cell-specific and sequence-based predictive models to these eQTLs to predict regulatory impacts and conducted heritability enrichment analyses, probabilistic colocalization, and transcriptome-wide association studies with GWAS of estimated glomerular filtration rate (eGFR) and urine albumin-to-creatinine ratio (UACR) (Teumer et al., 2019; Wuttke et al., 2019). A subset of candidate eQTLs were validated experimentally *in vitro* and using a *Drosophila* nephrocyte model. Altogether, we demonstrated improved precision in discerning putative functional SNPs within eSNP haploblocks, which subsequently increased discovery and provided biological insight in downstream analyses. Our interactive resource is available to the public at www.nephqtl2.org.

**Figure 1.**
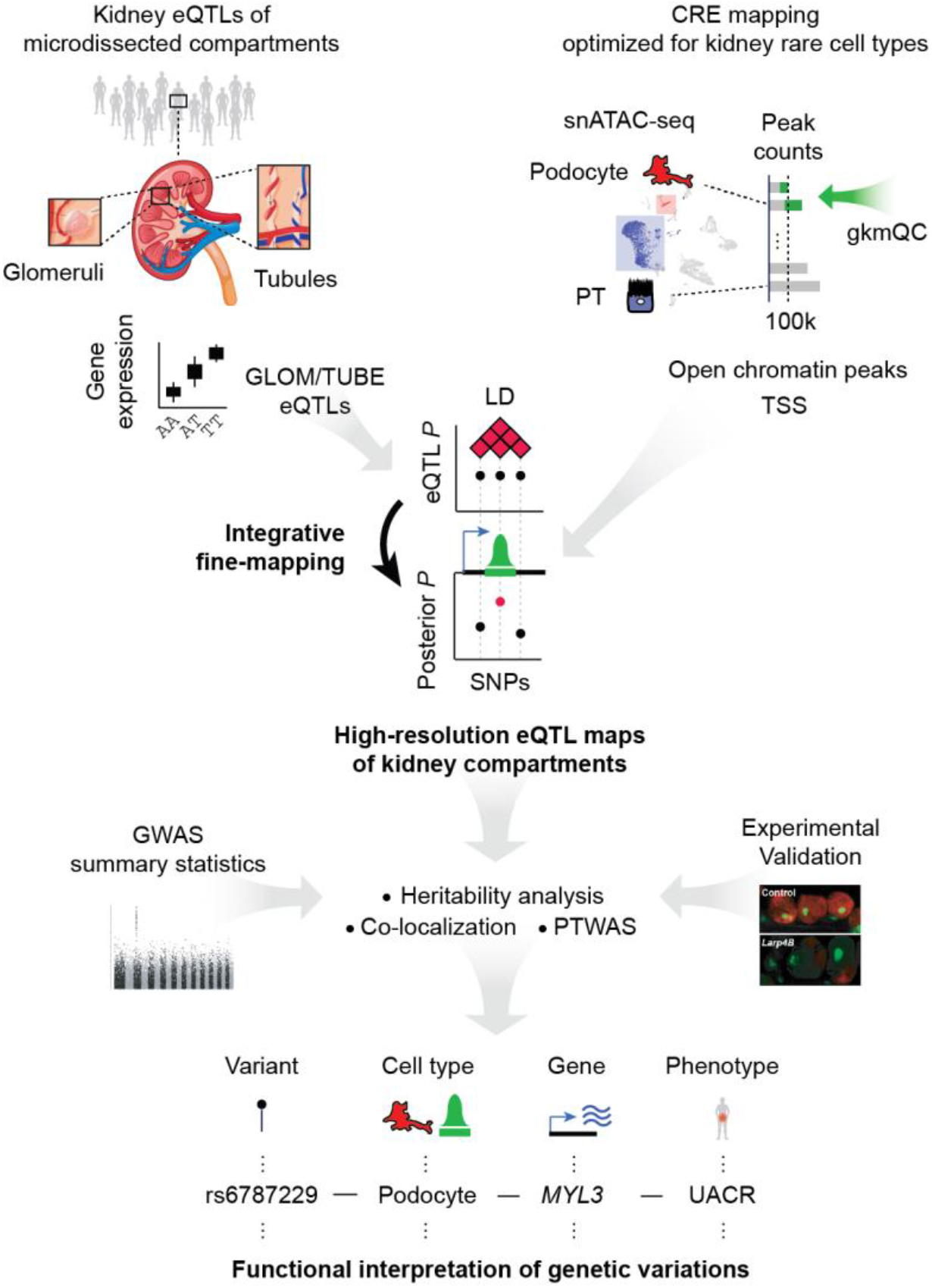
Analysis schematic: Integrating eQTLs with cell-type *cis*-regulatory annotations to build high-resolution eQTL maps of micro-dissected glomeruli (GLOM) and tubulointertisium (TUBE) and Downstream analyses for functional interpretation of genetic variations associated with kidney functional traits: eQTL = expression quantitative trait loci, CRE = *cis*-regulatory element, snATAC-seq = single nuclear assay for transposase-accessible chromatin using sequencing, gkmQC = gapped k-mer SVM quality check, TSS = transcription start site, LD = linkage disequilibrium, SNP = single nuclear polymorphism, GWAS = genome-wide association study, PTWAS = probabilistic transcriptome-wide association study, MYL3 = Myosin Light Chain 3, UACR = urine albumin-to-creatinine ratio.

## Results

### Multi-SNP fine-mapping of *cis*-eQTLs incorporating cell-type open chromatin annotations provides high-resolution eQTL maps

The eQTL analysis consisted of 332 NEPTUNE (Gadegbeku et al., 2013) individuals with paired RNA-seq and whole-genome sequencing (WGS) data, including 240 glomerular samples (GLOM) and 311 tubulointerstitial samples (TUBE; **Table S1**). For open chromatin annotations, we used kidney cell *cis*-regulatory element (CRE) maps that we recently created through the development and application of a new method to optimize the discovery of rare cell-type specific peaks using underlying sequence signatures - gkmQC (Han et al., 2022).

We first compared our optimized single-nucleus kidney CRE maps (Han et al., 2022) to bulk kidney data to show that the single-nucleus data detected 62% additional open chromatin regions not detected in bulk kidney data (Lee et al., 2022) (**Figure 2A)**. Beyond the increased quantity and uniqueness of CREs identified, several metrics indicated that our kidney single-nucleus CRE maps were of high quality. First, the statistical overlap of open chromatin peaks sorted the kidney cell types into four groups (which we denote as “C1” through “C4”) that reflected functional similarities and physical location in the nephron (**Figure 2B)**. By conducting stratified LD-score regression (S-LDSC) (Finucane et al., 2015) with cell-group-specific peaks, we found enriched heritability for UACR GWAS variants residing within C1-specific peaks (6.85-fold; *P* = 0.02), which includes peaks specific for podocytes and parietal epithelial cells. eGFR heritability was enriched in SNPs within C2 and C3-specific CREs, which include proximal tubule (*P* = 0.05) and loop of Henle (*P* = 0.04) specific peaks, respectively (**Figure 2C; Table S2)**. Additionally, the heritability enrichment for UACR increased with respect to groupings of peaks with higher podocyte and parietal epithelial cell specificity (**Figure 2D**).

**Figure 2.**
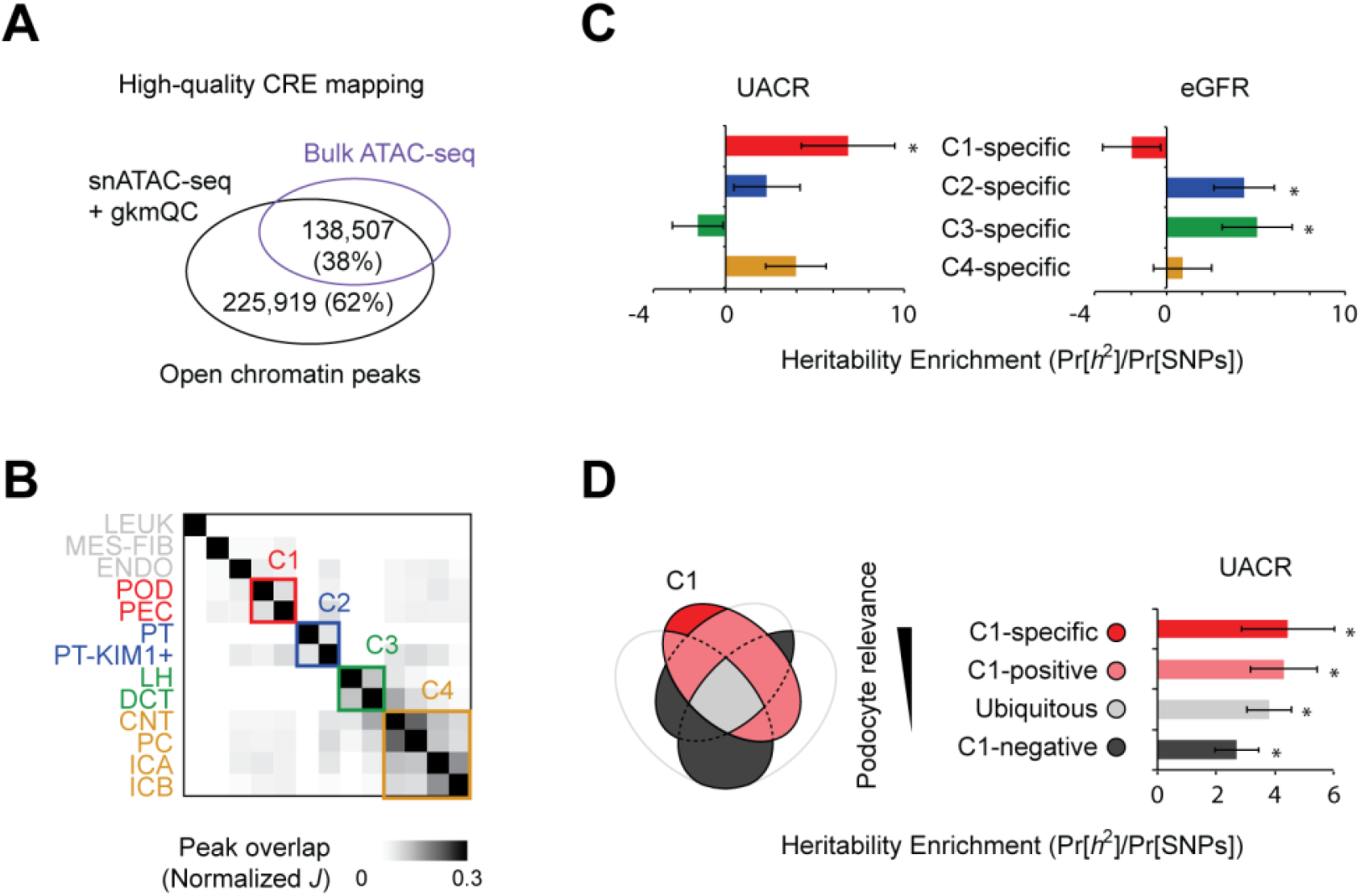
High-quality CREs and their cell-type specific contribution to the heritability of functional kidney traits. **(A)** Overlap between peaks of kidney snATAC-seq and bulk ATAC-seq from different samples: CRE = *cis*-regulatory element, snATAC-seq = single nuclear assay for transposase-accessible chromatin using sequencing **(B)** Heatmap presents the normalized Jaccard index (*J*) of the peak overlap between cell types used to form cell groups (**Methods and Materials**). C1-4 are groups that include kidney cell types from different kidney compartments; C1 = glomerulus, C2 = proximal tubule, C3 = the loop of Henle, C4 = collecting duct, LEUK = leukocytes, MES-FIB = mesangial/fibroblast, ENDO = endothelial, POD = podocyte, PEC = parietal epithelial, PT = proximal tubule, PT-KIM1+ = KIM1 positive proximal tubule, LH = loop of Henle, DCT, distal convoluted tubule, CNT = connecting tubule, PC = principal cells, ICA = type A intercalated cells, ICB = type B intercalated cells **(C)** The bar graph presents the heritability enrichment partitioned by the genomic coordinates of cell-group-specific open chromatin peaks; * *P* ≤ 0.05: UACR = urine albumin-to-creatinine ratio, eGFR = estimated glomerular filtration rate **(D)** The bar plot compares the heritability enrichments for urine albumin-to-creatinine ratio for the subgroups of peaks stratified by relevance to the C4 group (POD and PEC); * *P* ≤ 0.05

We used MatrixEQTL (Shabalin, 2012) for our single-SNP eQTL analysis, a necessary precursor to the ultimate multi-SNP fine-mapping step. The effect sizes from these results were comparable to those from eQTL studies of bulk kidney cortex (The GTEx Consortium, 2020) and of GLOM and TUBE from nephrotic syndrome (NS) (Gillies et al., 2018) and non-NS (Qiu et al., 2018) samples (**Figure S1**). A principal component analysis of eQTL *z*-scores from GLOM, TUBE, and GTEx tissues found that GLOM and TUBE were proximal to the kidney cortex and other non-brain tissues (**Figure S2A**). GLOM eSNPs showed the highest enrichment in podocyte and parietal epithelial cell open chromatin peaks, and TUBE eSNPs showed the highest enrichment in proximal tubule peaks (**Figure 3A-B**). Cell-type enrichment is also higher than randomly expected (**Figure S2B**). Of note, SNPs identified uniquely by our optimized peak calling method significantly contributed to cell-type enrichment (**Table S3**). Similarly, GLOM eGenes were most enriched for expression in podocyte and parietal epithelial cells, and the TUBE eGenes were most enriched for expression in proximal tubule cells (**Figure 3C**). Finally, in contrast to kidney cell types, no tissues from GTEx or ENCODE had a significant enrichment bias towards either GLOM or TUBE eSNPs (Davis et al., 2018; The ENCODE Project Consortium, 2012) (**Figures S2C-D**). This indicates that the kidney single-nucleus-based approach is necessary to create functional annotation priors discriminating transcriptional landscapes of GLOM and TUBE eQTLs.

**Figure 3.**
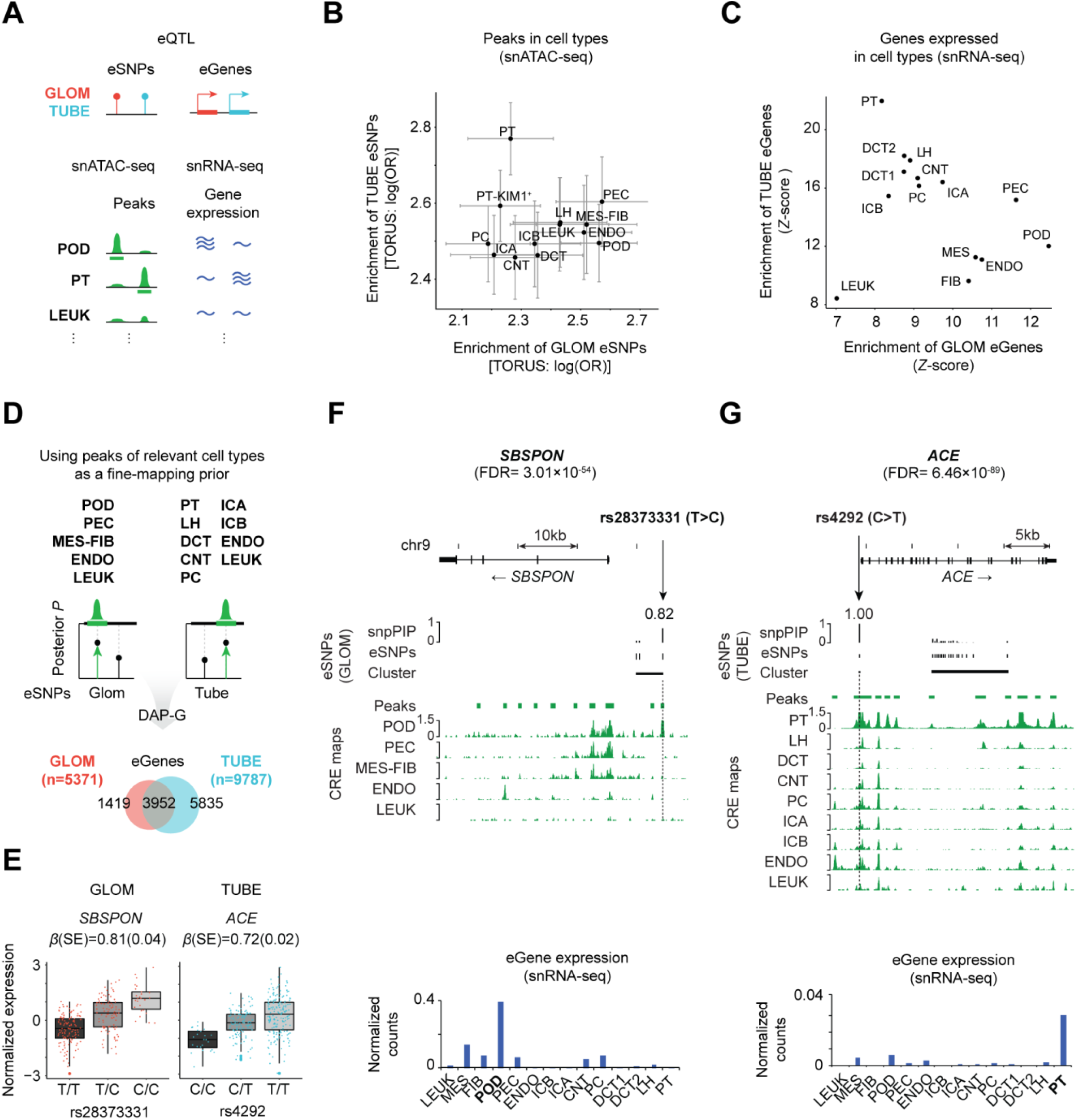
Integrative analysis of eQTLs and high-quality CRE maps. **(A)** A schematic demonstrates the cell-type-specific enrichment of eSNPs and eGenes for the peaks and genes from snATAC/RNA-seq datasets: eQTL = expression quantitative trait loci, eSNPs = single nucleotide polymorphisms associated with gene expression, eGenes = genes with at least one variant associated with expression, GLOM = glomerular eQTLs, TUBE = tubulointerstitial eQTLs, snATAC-seq = single nuclear assay for transposase-accessible chromatin using sequencing, snRNA-seq = RNA sequencing **(B)** Enrichment analyses of eSNPs in open chromatin peaks in corresponding cell types: log(OR) = natural logarithm of the odds ratio, LEUK = leukocytes, MES-FIB = mesangial/fibroblast, ENDO = endothelial, POD = podocyte, PEC = parietal epithelial, PT = proximal tubule, PT-KIM1+ = KIM1 positive proximal tubule, LH = loop of Henle, DCT, distal convoluted tubule, CNT = connecting tubule, PC = principal cells, ICA = type A intercalated cells, ICB = type B intercalated cells **(C)** Enrichment analysis of eGenes among all genes expressed in corresponding cell types. **(D)** Cell types used for CRE fine-mapping annotations for integration into DAP-G and resulting eGenes. DAP-G = deterministic approximation of posteriors, snpPIP = SNP posterior inclusion probability **(E)** Plots of the top ranked GLOM/TUBE-specific eQTLs. Coordinates are based on the hg19 build: *SBSPON* = Somatomedin B And Thrombospondin Type 1 Domain Containing, *ACE* = Angiotensin 1 Converting Enzyme, β (SE) = Effect size of genotype on gene expression and standard error from single-SNP association **(F-G)** Specific examples of fine-mapped eSNPs in the clusters of top GLOM/TUBE-specific eGenes. The heights of vertical black bars depict the snpPIP of each clustered eSNP. Horizontal black bars depict the genomic range of each cluster. Green horizontal bars depict the range of open chromatin peaks of the relevant cell types. The green vertical graph shows the normalized pile-up of snATAC-seq reads of the corresponding cell type. Blue bar plots present the gene expression of snRNA-seq data normalized by genes and cell counts.

We generated SNP priors by including our cell-type-specific open chromatin annotations and the distance between each SNP and the corresponding gene’s transcription start site (TSS). We refer to this as the “integrative prior” (**Methods and Materials)**. For our GLOM CRE annotation, all SNPs within podocyte, parietal epithelial, endothelial, mesangial/fibroblast and leukocyte open chromatin peaks were combined. For TUBE, we combined CRE from all proximal tubule clusters, loop of Henle, distal convoluted tubule, connecting tubule, principal cells, type A and B intercalated cells, endothelial cells, and leukocytes (**Figure 3D)**. We determined the weights for each SNP by integrating the estimated enrichment of our single-SNP eQTL associations among CREs into SNP priors with “TORUS” (Wen, 2016). Variants in open chromatin are 3.95 [CI: 3.76, 4.15] fold more likely to be eSNPs in the TUBE and 4.03 [CI: 3.72, 4.35] fold more likely in the GLOM.

The integrative priors were then used with our expression and WGS data to fine-map *cis*-eQTLs allowing for multiple independent SNPs per gene using the method “DAP-G” (Lee et al., 2018b; Wen et al., 2016). After Bayesian FDR control, we identified 5,371 GLOM and 9,787 TUBE eGenes (**Figure 3D, Table S4**). This is an approximately 6-fold increase in eGenes compared to our previous array-based analysis (**Figure S3**). These kidney CRE-informed eQTLs provided higher resolution in and of themselves. But using our single-nucleus RNA-seq (snRNA-seq) and snATAC-seq data to examine the eQTLs in a kidney cell-type-specific manner provided even greater resolution.

Our increase in resolution and interpretability is illustrated with the Somatomedin B And Thrombospondin Type 1 Domain Containing gene (*SBSPON*), the most significant GLOM-specific eGene (FDR = 3.01×10^−54^; **Figure 3E and F**). Before fine mapping, we found three eSNPs within the single associated haploblock with indistinguishable effects on *SBSPON* expression (**Table S5**). However, our multi-SNP fine mapping identified rs2837331 as the putative causal eSNP (SNP posterior inclusion probability (snpPIP) = 0.82). Inspection of the snATAC-seq data identified that this SNP was in a podocyte-unique open chromatin peak ~10kb upstream of the *SBSPON* locus. This is concordant with the snRNA-seq analysis showing its podocyte-specific expression (**Figure 3F)**.

Angiotensin I Converting Enzyme (*ACE*) is a TUBE-specific eGene with one of the top-ranked signals (FDR = 6.46×10^−89^; **Figure 3E and G**). Our fine mapping also found a putatively causal eSNP (rs4292; snpPIP = 1.00) in the *ACE* promoter that is open across multiple cell types, with the strongest peak in proximal tubules, which is concordant with its gene expression pattern. Taken together, our results suggest that this cell-type CRE-informed fine-mapping approach provides higher resolution eQTL maps, which improves our ability to dissect the transcriptional regulation in kidney tissues.

### CRE-informed eQTLs provide higher-resolution fine mapping and enriched heritability for kidney phenotypes

We next conducted a series of complementary genome-wide analyses to assess potential improvements in eQTL fine-mapping resolution that result from using an integrative prior versus a (1) “uniform prior” - all SNPs having equal prior probability of being an eSNP, and (2) “TSS prior” - only including the distance from the TSS.

The first metric was the posterior probability of each cluster’s lead SNP (snpPIP), where an increase would indicate more confidence being shifted to the lead SNPs. The second metric was the number of SNPs in the eQTL’s 95% credible sets, where a decrease would indicate an improvement. On both metrics, the integrative prior showed the highest fine-mapping resolution, followed by the TSS prior alone, then a uniform prior (**Figure 4A; Figure S4A**).

**Figure 4.**
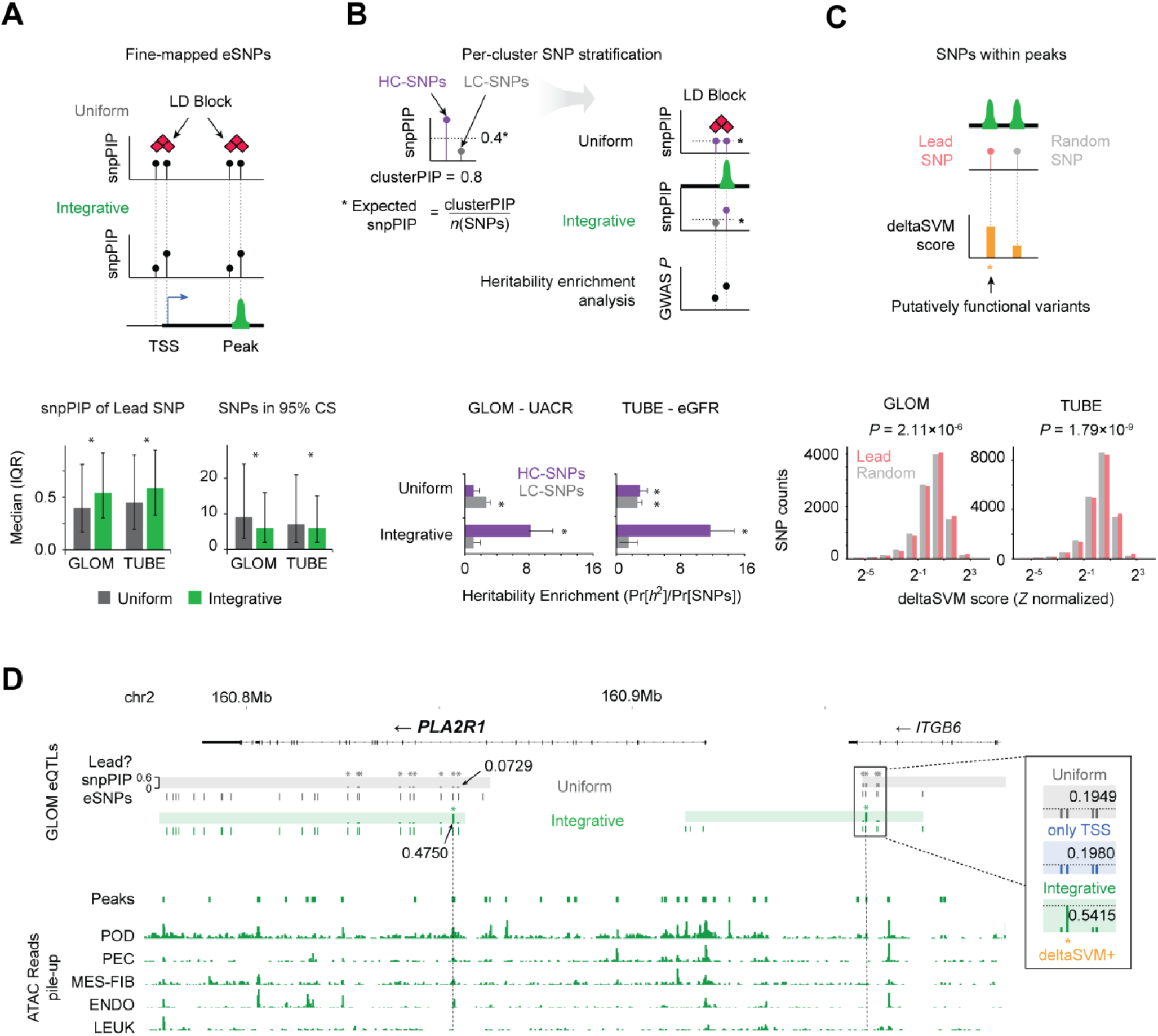
Specifying putative functional variants using the high-resolution eQTL map. **(A)** A schematic showing how fine mapping with an integrative prior stratifies putative functional variants among eSNPs within the same LD block (top). For each eGene cluster in GLOM and TUBE, we compare the distribution of the top SNPs’ snpPIPs and the number of SNPs forming the 95% credible set between the uniform and integrative priors (bottom). The distributions are significantly different for all four comparisons, Wilcoxon test *P* < 2×10^−16^; eSNPs= single nucleotide polymorphisms associated with gene expression, LD = linkage disequilibrium, snpPIP = SNP posterior inclusion probability, TSS = transcription start site, CS = credible set, IQR = interquartile range **(B)** A schematic demonstrates how high/low-confident eSNPs (HC/LC-SNPs) are stratified per cluster and how the integrative prior weighs the putative functional SNPs enriched for the heritability of relevant traits. Bottom bar plots compare the heritability enrichment of GLOM/TUBE HC/LC-SNPs fine mapped with the two different priors (uniform and integrative). GWAS traits and eQTLs are paired based on the cell-type convergence of the traits derived from our heritability enrichment analysis (**Figure 2E**) and the *a priori* knowledge of the trait physiology. Asterisks depict the significant enrichment of the heritability assessed by S-LDSC *P* ≤ 0.05: LD = linkage disequilibrium, snpPIP = SNP posterior inclusion probability, GWAS = genome-wide association study, UACR = urine albumin-to-creatinine ratio, eGFR = estimated glomerular filtration rate **(C)** A schematic diagram demonstrating the hypothesis that high-resolution eQTLs are more likely to be functional— inferred by the deltaSVM model—than random common SNPs within open chromatin peaks. Random SNPs have similar genomic properties with high-resolution eQTLs. Statistical significance of the difference between deltaSVM scores is assessed by the Wilcoxon rank-sum test. **(D)** The genomic coordinate diagram presents a detailed comparison between the fine mapping of eSNPs with the integrative and uniform priors. Phospholipase A2 Receptor 1 (*PLA2R1*) and the change of lead SNPs and PIPs across priors for cluster 1 and cluster 3, are presented. For the integrative prior, the top SNPs are rs73967997 for cluster 1 and rs12987602 for cluster 3. Gray and green bands are the ranges of clusters from uniform and integrative priors, respectively. Asterisks mark the lead SNP per cluster and arrows indicate snpPIP scores. Vertical dashed lines connect the lead SNPs to their genomic coordinate with an open-chromatin perspective. Four lead SNPs of interest from the right cluster are presented in the sub-panel, where snpPIPs from the TSS-only prior (blue) and deltaSVM positive variants (orange) are also shown. Bottom: Horizontal and vertical bar plots depict the genomic ranges of open chromatin peaks and the pile-up of snATAC-seq reads for relevant cell types: LEUK = leukocytes, MES-FIB = mesangial/fibroblast, ENDO = endothelial, POD = podocyte, PEC = parietal epithelial, *ITGB6* = Integrin Subunit Beta 1.

The third metric was a change in S-LDSC-based heritability enrichment of “high confidence” (HC) versus “low confidence” (LC) eSNPs as a function of prior choice. We defined HC-eSNPs as those whose posterior inclusion probability (PIP) within a haploblock cluster is higher than would be observed if the PIPs were equally distributed among all SNPs in a locus. LC-eSNPs have snpPIPs smaller than expected (**Figure 4B**). There was a significant enrichment of heritability for HC-SNPs using the integrative prior for UACR and eGFR; this was greater than the heritability observed in analyses using the uniform and TSS-only priors (**Figure S4B**).

The final metric was a change in the computationally-predicted functionality of fine-mapped eSNPs defined using the different priors. To do this, we compared the deltaSVM score (Lee et al., 2015) of lead eSNPs to random SNPs in the CREs, controlling for allelic frequency, distance from TSS, and the signal strength of open chromatin peaks. Lead eSNPs from the integrative prior had significantly higher deltaSVM scores than the random SNPs (*P* = 2.22×10^−9^ for GLOM; *P* = 4.64×10^−6^ for TUBE) (**Figure 4C**) and those fine-mapped by uniform priors (*P* = 5.65×10^−7^ for GLOM; *P* = 1.49×10^−17^ for TUBE) (**Figure S4C**).

A focused analysis of *PLA2R1* illustrates the power of these high-resolution eQTL maps to more accurately specify the putative functional variants. *PLA2R1* is a glomerular gene specifically expressed in podocytes (**Figure S4D**) that is associated with a rare kidney disease, membranous glomerulonephritis (Beck et al., 2009). Our fine mapping identified four independent eSNP clusters in GLOM. We highlight two clusters in **Figure 4D** where the lead eSNPs identified using the integrative prior have a higher posterior probability than those identified using the uniform prior. The lead SNPs from the integrative prior were in podocyte-specific open chromatin peaks, concordant with gene expression pattern of *PLA2R1*, and are deltaSVM positive, implying that the lead SNPs newly found using the integrative prior are more likely to be causal variants.

### Colocalization of high-resolution eQTLs and kidney-relevant GWAS SNPs identifies novel genes and increased resolution of colocalized signals

Including cell-type-informed CREs as integrative priors in our eQTL analyses led to increased posterior SNP probabilities of putative regulatory SNPs and, in some cases, distinguished between SNPs in high LD. Given this, we hypothesized that colocalization analysis with well-powered GWAS of eGFR and UACR and these high-resolution eQTL maps would increase detection and fine-mapping resolution of colocalized signals. To identify colocalized SNPs, we used fast enrichment estimation aided colocalization analysis (fastENLOC) (Pividori et al., 2020; Wen et al., 2017). For eGFR, we identified 46 TUBE and 6 GLOM colocalization signals, which we defined as having a regional colocalization probability (RCP) ≥ 0.5. For UACR, there were 9 TUBE and 21 GLOM colocalization signals (**Table S6, Figure 5A; Figure S5A-B**). When comparing colocalization signals for kidney traits derived from previous array-based kidney eQTLs (Teumer et al., 2019; Wuttke et al., 2019), we replicated five genes for UACR – *PRKCI, TGFB1, PTH1R, MUC1, OAF* – and 3 for eGFR – *FGF5, MLLT3, UMOD*. Using the high-resolution eQTLs, we discovered 82 colocalized loci, with 22 of them fine mapped to a single variant. 90% of these single variants (18/22) were in open chromatin. In contrast, we only identified 69 colocalized loci with the uniform prior. Finally, from a systematic comparison of extended sets of colocalized SNPs (RCP ≥ 0.2) across the different priors, we confirmed a significant increase in colocalization probability when high-resolution eQTLs are incorporated (**Figure S6**).

**Figure 5.**
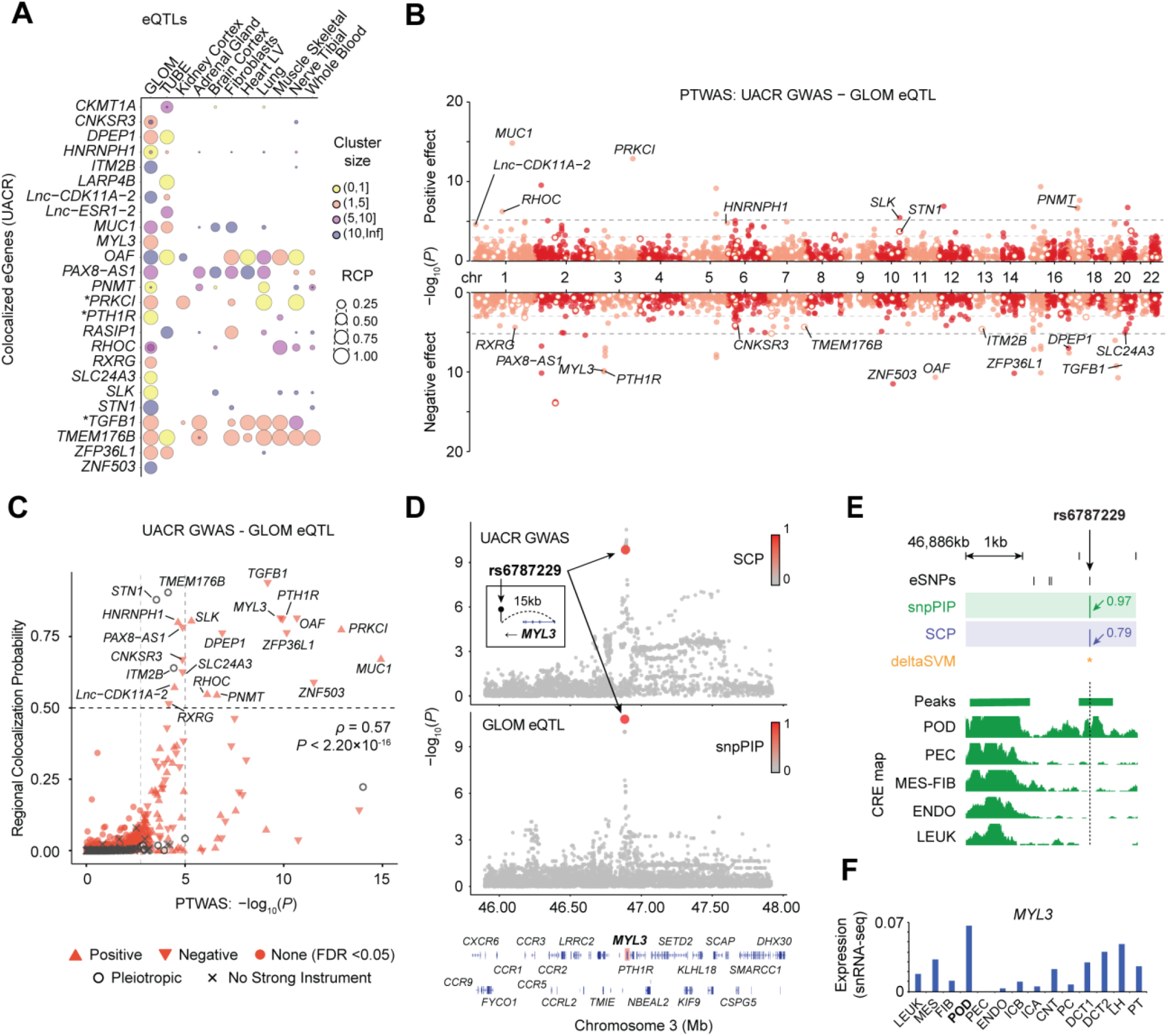
PTWAS and colocalization analyses of GLOM high-resolution eQTLs and UACR GWAS. **(A)** A bubble plot shows GLOM or TUBE eGenes colocalized with UACR GWAS loci (≥1 cluster with Regional colocalization probability (RCP) ≥ 0.5) along with GTEx results from select tissues, including kidney cortex. Each circle represents an eQTL cluster, and its diameter and color depict the RCP scores and the cluster size (number of eSNPs), respectively; *gene replicated from previous analyses, all other genes are novel. **(B)** A Miami plot shows the PTWAS signal of putative causal genes associated with the variance of UACR measured by transcriptional changes imputed by high-resolution eQTL. Genes lacking a strong instrument are excluded from this figure. Each dot represents the gene, and the genes potentially confounded by pleiotropic effects are indicated with an open circle. Dashed lines indicate thresholds using two multiple testing correction methods, *q*-value (*q* ≤ 0.05; light gray) and Bonferroni (*P* ≤ 9.52×10^−6^; dark gray). Glomerular eGenes in the bubble plot of (A) are annotated. **(C)** A scatter plot of RCP of top colocalized clusters from fastENLOC and corresponding PTWAS associations for each eGene. The shape of points depicts the type of effect of the PTWAS association. **(D)** Summary plots of GWAS and eQTL show SNP associations from the UACR GWAS and GLOM eSNPs in the *cis*-window (≤1Mb from the gene body) of *MYL3*. GWAS hits and eSNPs are colored by SNP-level colocalization probability (SCP) from fastENLOC and snpPIP from eQTL fine-mapping, respectively. **(E)** Top: Four eSNPs in the top fastENLOC cluster of *MYL3* are displayed. Green and purple vertical bar plots depict the snpPIP and SCP, respectively. Orange asterisk depicts the putative-functional variants inferred by the deltaSVM model. Bottom: Green horizontal and vertical bar plots depict the genomic range of open chromatin peaks and the pile up of snATAC-seq reads on the relevant cell types. **(F)** Blue bar plots present the normalized expression of *MYL3* in snRNA-seq data

Previous studies have shown that functional GWAS variants are enriched in CREs of relevant tissues and cell types (Maurano et al., 2012). Thus, we hypothesized that if a specific prior better captured the functional GWAS variants, then lead colocalized SNPs weighted by that prior would be enriched in CREs more than expected. Indeed, we found that colocalized SNPs found using our high-resolution eQTLs were significantly enriched in CREs, while colocalized SNPs exclusively from our uniform prior were not **(Table 1)**. Our results suggest that the CRE-informed colocalization analysis promotes the discovery of the functional GWAS variants.

These high-confidence potential target genes and SNPs discovered by our colocalization analyses can be a starting point for mapping GWAS variants to their function. For example, the replication of *PTH1R* highlights the increased resolution and cell-type interpretation. Compared to Teumer et al., we refined the *PTH1R* eSNP credible set size from 14 to 1. The lead SNP also changed from rs73065147 (SNP colocalization probability (SCP) = 0.2), which does not fall within any CRE, to rs6787229 (SCP = 0.79), which resides in a podocyte-specific peak (**Figure S7)**. Thus, by weighing eSNPs within CREs, we more confidently identified putative causal SNPs and hypothesized an association between podocyte-specific regulation of *PTH1R* and UACR.

### Probabilistic transcriptome-wide association analysis (PTWAS) with high-resolution eQTLs identifies associations between SNP-predicted gene expression and kidney phenotypes

Using our high-resolution eQTL maps for the predictive model of gene expression, we next analyzed the association between SNP-predicted gene expression and eGFR and UACR using probabilistic transcriptome-wide association study (PTWAS) (Zhang et al., 2020) (**Methods**). For eGFR, at a false discovery rate (FDR) ≤ 5%, we identified 601 significant gene-trait pairs in GLOM and 1,074 in TUBE. For UACR at an FDR ≤ 5%, we identified 137 significant gene-trait pairs in GLOM (**Figure 5B**) and 179 in TUBE **(Table S7, Figures S8A-C)**. We also found a significant correlation (*ρ* = 0.57, *P* ≤ 2.2×10^−16^) between colocalization and PTWAS signals (**Figure 5C, Figure S8D-F**), demonstrating the consistency of inference results when different analytical approaches are applied to the same dataset (Hukku et al., 2021a).

Our integrative analysis enables us to interpret cell types and CREs in which GWAS variants regulate their target genes. As a representative case, we highlight a colocalized SNP, rs6787229, associated with *MYL3* gene expression and UACR also validated by PTWAS (*P* = 1.44×10^−10^, SCP = 0.79; **Figure 5D**). A podocyte-specific open chromatin peak harboring this SNP increased probability of the eQTL fine-mapping (snpPIP = 0.97) compared to fine mapping with the uniform prior (snpPIP = 0.81) (**Figure 5E**). This inferred cell-type specificity was corroborated by podocyte-specific gene expression of target gene *MYL3* (**Figure 5F**). Taken together, these integrative approaches with high-resolution eQTL maps increase the opportunity to map variants to function via gene regulation with greater interpretability and confidence.

### SNP- and gene-level validation of predicted-causal eQTLs results in reduced *Drosophila* nephrocyte function and SNP-level regulation of *LARP4B* and *NCOA7*

We identified GLOM and TUBE eGenes that were (1) significant in both colocalization and PTWAS analyses with UACR and/or eGFR, (2) contain colocalized SNPs in CREs, and (3) had gene homolog expression in *Drosophila* nephrocytes. Fourteen of these 32 resulting genes were randomly selected for experimental validation **(Table S8; Methods and Materials)**. To do this, we used an *in vivo Drosophila* model. The *Drosophila* nephrocytes filter and reabsorb circulating proteins in the hemolymph and share many similarities with glomeruli and tubule cells at the functional, molecular, and ultrastructural levels (Weavers et al., 2009; Zhang et al., 2013a), making it an ideal model for both GLOM and TUBE eGenes. In flies carrying MHC-ANF-RFP transgene, the myosin heavy chain (MHC) promoter directs muscle cell expression of a rat atrium natriuretic factor (ANF)–red fluorescent protein (RFP) fusion protein (ANF-RFP) that is secreted into the hemolymph (Zhang et al., 2013b). ANF-RFP is typically filtered and endocytosed by healthy, wild-type nephrocytes, and the intracellular red fluorescence can be readily visualized and quantitated *in vivo*. We found nephrocyte-specific knockdown of five genes impacted nephrocyte function -- *Fkbp12* (*FKBP1A*), *Larp4B* (*LARP4B*), *Mlc-c* (*MYL3*), *mtd* (*NCOA7*), and *svr* (*CPXM1*) (**Figure 6**). In an independent *ex vivo* functional assay, we tested the ability of dissected nephrocytes to absorb Texas Red-labeled 10 kD Dextran particles. Consistent with the secreted protein reabsorption assay, silencing anyone of the five genes resulted in a decrease in intracellular Texas Red fluorescence compared to control nephrocytes. Of note, six genes tested in this experiment (*ZFP36L1, NDRG1, GLUD1, XPC, HLF*, and *CG3662*) were assessed as pleiotropic in the PTWAS and were not found to impact nephrocyte function when knocked down. To further validate TUBE associations identified in *LARP4B* and *NCOA7*, we generated luciferase reporter constructs **(Table S9)** to test the allele-specific enhancer activity of the lead variants associated with each gene, rs80282103 and rs11154336, respectively. Consistent with *LARP4B* eQTL findings, the rs80282103-T minor allele demonstrated 126% increased reporter activity in HK-2 cells, a human proximal tubule cell line (*P* = 5.99 × 10^−9^), in both “forward” and “reverse” orientations. The rs11154336-A allele demonstrated 236% increased reporter activity compared to the G allele, consistent with *NCOA7* eQTL results (*P* = 5.99 × 10^−9^) **(Figure S9)**.

**Figure 6.**
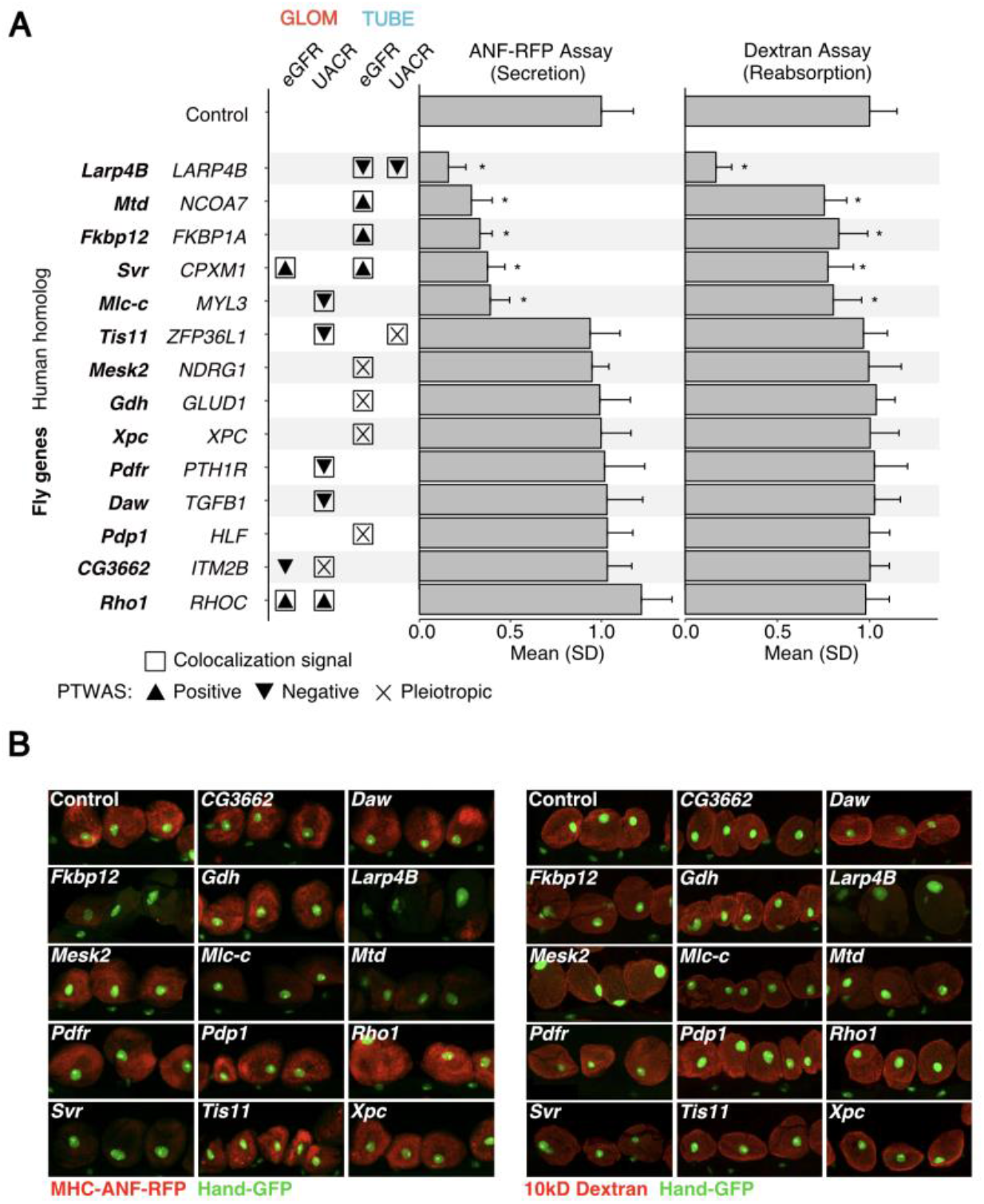
The impact on *Drosophila* nephrocyte function caused by RNAi of the kidney trait-associated genes. **(A)** Comparison of 14 genes and RNAi control. *Drosophila* genes are followed by human homologs and association indicator from tissue-phenotype analyses; colocalization (RCP ≥ 0.5) and PTWAS associations (FDR ≤ 0.05), including direction of effect. Bar plots indicate mean fluorescence of 60 nephrocytes (20 nephrocytes from three flies). Significant differences (t-test) between the gene and control are indicated (* *P* ≤ 0.001). **(B)** Fluorescence microscopy images for representative nephrocytes for each *Drosophila* gene. Left panel: MHC-ANF-RFP assay; Right panel: 10kD Dextran; Hand-GFP = nephrocyte nucleus.

## Discussion

In mapping the non-Mendelian genomic basis of kidney traits and diseases, we are challenged to maximize the detection of regulatory circuits - functional genetic variants in *cis*-regulatory elements, their target genes, and their cells of action. Conventional fine-mapping approaches, depending on the population size and haplotype structure, may be suboptimal in specifying the putative functional variants with low allelic frequency or multiple indistinguishable tag SNPs within the same haplotype block. Given that the functional characteristics of variants in CREs are orthogonal to LD patterns, diverse functional annotations have been used for fine mapping of GWAS and eQTL variants. To this point, we created a workflow that used single-nucleus open chromatin data to generate priors for use in Bayesian multi-SNP eQTL detection algorithms. In doing so, we demonstrated improved precision in discerning putative functional SNPs within eSNP haploblocks (“fine-mapping”), which subsequently increased discovery and biologic insight of downstream analyses.

The CRE-informed fine mapping for eQTLs can recover underpowered variants with a Bayesian approach that augments the enrichment of eQTLs across tissue-relevant cell-type CREs, reflecting the underlying biology of transcriptional regulation. Corroborating this, we found that the gain in heritability and deltaSVM score enrichment, when comparing our integrative approach to results from a uniform prior, was larger in the lower-powered tissue, GLOM, compared to our higher-powered TUBE analysis (**Figure 4**). In sum, these integrative approaches can complement the mapping of eQTLs with moderate statistical power and be an effective resource for the discovery of eQTLs from the limited samples with nephrotic syndrome.

A particularly important aspect of this study was our ability to use our newly developed method, gkmQC, to characterize CREs in rare cell types at a coverage and level of resolution not previously attained. For example, it was critical to comprehensively map (putatively)-casual regulatory variants of podocytes, a key rare (<1%) cell type involved in kidney filtration function. By nearly doubling the podocyte-specific peaks, we increased the statistical power of enrichment between podocyte peaks and glomerular eQTLs, which in turn improved fine-mapping efforts. This directly led to our ability to discover, via LD score regression analysis, a significant contribution of podocyte regulatory elements to the heritability of urine albumin-to-creatinine ratio (UACR) from GWAS studies. Interestingly, we also found enrichment of eGFR-associated SNPs among proximal tubule, loop of Henle, and distal convoluted tubule open chromatin. Together, these underscore the ability of our optimized CRE maps to provide biological insights into multiple kidney phenotypes and should serve as a resource to investigators seeking to discern the specific regulatory circuits of these diseases and traits.

As intended, our integrative approach allowed us to increase fine-mapping resolution, marked by smaller credible set sizes and increased statistical confidence of lead SNPs. To highlight the utility of our eQTLs in mapping variants to their function, we performed colocalization and transcriptome-wide association studies with functional kidney outcomes UACR and eGFR. Colocalization analyses are hindered by low power, especially when LD matrices are not perfectly matched (Hukku et al., 2021b). We found that weighing putative functional SNPs in our eQTL analysis resulted in an enrichment of colocalized SNPs within CREs and increased discovery of novel loci, thus partially overcoming this power limitation. These complementary analyses not only highlight the utility of our eQTL resource but also allow for new biological insights into associations between tissue and cell-specific gene regulation and kidney function. This was illustrated by a high confidence eSNP within a podocyte-specific CRE that was associated with both *MYL3* glomerular expression and UACR.

By following up on statistical findings in the *Drosophila* nephrocyte, we were able to further validate selected genes of interest. For example, we replicated the association between a single intronic *LARP4B* eSNP, rs80282103, and both UACR and eGFR, previously discovered by Wuttke et al. and Morris et al. (Morris et al., 2019; Wuttke et al., 2019) and identified a novel association between an intronic *NCOA7* eSNP rs11154336 and eGFR. We found that knockdown of *Larp4b* and *Mtd*, an *NCOA7* ortholog, in the *Drosophila* nephrocyte had the most statistically significant reductions in nephrocyte function, providing orthogonal support for the functional role of *LARP4B* and *NCOA7* in the kidney. Using a luciferase assay, we also validated the functional impact of rs80282103 and rs11154336. In addition to *LARP4B* and *NCOA7*, three novel colocalized genes —*FKBP1A, CPXM1*, and *MYL3* — impacted secretion and reabsorption by the nephrocyte. In our colocalization analysis, we identified a single SNP associated with both *MYL3* and *PTH1R* expression and UACR. Interestingly, only *MYL3* knockdown impacted nephrocyte function, providing support for the role of *MYL3* (vs. *PTH1R*) in kidney function. We selected follow-up genes independent of their predicted horizontal pleiotropic effects. Interestingly, we found that all six genes predicted to be pleiotropic did not impact *Drosophila* nephrocyte function **(Figure 6)**.

The eQTLs augmented by cell-type CREs make the results of downstream analyses (colocalization, PTWAS) interpretable in terms of (1) mechanistic insight into transcriptional regulation and (2) contributing cell-types or *cis-*regulatory elements (**Figure 5-6**). The clinical implication of such interpretable eQTLs can be inspected by the researchers with reduced false-positive hits arising from neutral variants in LD with causal variants. To facilitate secondary analyses for end users, we provide interactive visualizations that include eQTL summary statistics along with cell-type CREs and deltaSVM scores at www.nephqtl2.org. This portal will be a novel resource to narrow down potential mechanisms and elucidate the regulatory landscape of kidney phenotypes.

### Limitations of study

The current study is limited in several ways: (1) Although we maximized the discovery of open chromatin peaks of rare cell types, the capability of peak discovery is still limited by the mappability of ATAC-seq reads, which depends on absolute cell counts. By harmonizing this data with data from future assays, we will be able to increase the cell counts and enhance the sensitivity of peak calling. (2) fastENLOC analysis, as well as other colocalization methods, tend to yield few highly confident findings. This is partially because the GWAS and the eQTL data are from different cohorts (the two-sample design), and their LD patterns do not match exactly. In addition, when working with summary statistics, an LD matrix from a third orthogonal population is used, which may not perfectly match the GWAS and eQTL datasets. However, while LD pattern mismatches reduce enrichment estimates and power, false positives are rare (Hukku et al., 2021b). Additionally, the comparison of colocalized loci with previous studies is imperfect due to the use of different methods. (3) Our eQTL dataset was built from a heterogenous nephrotic syndrome cohort from multiple ancestries. While this may allow us to improve fine-mapping (Wen et al., 2015) and capture disease-specific eQTLs, the heterogeneity had to be properly controlled. To this end, we adjusted our eQTL analysis with PEER factors, which account for hidden technical and biological structure, and principal components, which account for population stratification. Interpretation of the eQTLs should take these factors into account. Of note, we found strong concordance of our eQTL effect sizes with compartment-matched eQTLs from healthy European samples (Qiu et al., 2018) **(Figure S1)**.

## METHODS AND MATERIALS

### Analysis of snATAC/snRNA-seq data

We used our optimized kidney CRE maps generated from a previous study (Han et al., 2022) using publicly available human kidney snATAC-seq data from non-tumor kidney cortex samples from 5 patients undergoing partial or radical nephrectomy (GSE151302) (Muto et al., 2021). Briefly, our optimized pipeline for snATAC-seq data processing includes (1) quality control of reads and cells with the Cell Ranger ATAC pipeline (v1.1.0), (2) harmonizing samples (Korsunsky et al., 2019), (3) cell clustering and type identification with snapATAC (Fang et al., 2021), (4) *post hoc* peak calling and optimization in a cell-type resolved manner with MACS2 (Zhang et al., 2008) and gkmQC (Han et al., 2022). A total of 35,286 cells were analyzed to call the peaks. Regarding snRNA-seq, we downloaded count matrices and cell-type labels for the five snRNA-seq samples (GSE151302) to measure gene expression and the cell type identification of snATAC-seq datasets. Consequently, we profiled open chromatin peaks and gene expression for 16 known kidney cell types.

### Analysis of bulk ATAC- and DNase-seq data

For the kidney bulk ATAC-seq data, we used the processed data obtained from Dr. Chakravarti’s laboratory (Lee et al., 2022). We obtained representative ENCODE DNase-seq samples of seven bulk tissues (Davis et al., 2018) with the best quality chosen by gkmQC. To process bulk DNase-seq and ATAC-seq, we adapted the previously established framework for DNase-seq and bulk ATAC-seq analyses as used in Lee et al. (Lee et al., 2018a) and Nandakumar et al. (Nandakumar et al., 2020). We used the same *post hoc* peak calling and optimization as used in snATAC-seq analysis gkmQC (Han et al., 2022).

### Heritability enrichment analysis

We used stratified LD-score regression (S-LDSC) (Finucane et al., 2015) to estimate the proportion and enrichment of heritability from GWAS summary statistics. The proportion of the heritability contributed by a SNP set (*C*) in open chromatin peaks from a sample is calculated as follows:

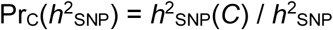

The enrichment of proportional heritability then is calculated by Pr_*C*_(*h*^2^)/Pr_*C*_(*M*), where Pr_*C*_(*M*) is the proportion of SNPs in *C* among the total SNP set (M). For reference LD scores, European ancestry population and the corresponding allele frequencies in 1000 Genomes Phase 3 data were used (v2.2; https://data.broadinstitute.org/alkesgroup/LDSCORE/). We processed open chromatin regions following our previously established steps (Han et al., 2022) to include the set of potentially associated regulatory variants. When comparing multiple functional annotations (e.g., multiple groups of CREs specific for different cell types), we conducted S-LDSC regression jointly with the annotations, along with the full set of the baseline annotations.

### Kidney RNA-seq

Total RNA from microdissected biopsies (240 GLOM, 311 TUBE) from the NEPTUNE study (Gadegbeku et al., 2013) were prepared using the Clontech SMARTSeq v4 kit. Samples underwent sequencing using Illumina HiSeq 2500, resulting in 150bp unstranded, paired-end reads. Fastq files underwent quality control filtering and trimming using fastQC, fastQScreen (Wingett and Andrews, 2018), and Picard Tools (http://broadinstitute.github.io/picard). Trimmed reads were aligned to the human genome (GRCh37) with STAR 2.6.0a (Dobin et al., 2013). Gene expression counts were quantified using StringTie v2.1.4 (Pertea et al., 2015). Gene expression was normalized across samples using TMM normalization with edgeR (Robinson et al., 2010), only keeping genes with greater than 0.1 cpm (~ 6 counts) in 20% of the samples. Transformed expression values were then rank-based inverse normalized.

### WGS

Whole genome sequencing (30x) was done using the Illumina HiSeq system. Alignment and variant calling were performed using default settings of GotCloud with the GrCh37 reference of the human genome (Jun et al., 2015). Variants underwent the following quality control filters using VCFtools (Danecek et al., 2011), PLINK (Purcell et al., 2007) and the HardyWeinberg R(v3.5.1) package (Graffelman, 2015): multi-allelic variants were converted to bi-allelic, variants with GQ < 20 and AB < 0.2 or > 0.8 were set to missing, variants with genotyping rate < 0.85, MAF < 0.01 and inbreeding coefficient < −0.3 were removed, and variants failing HWE, *p* < 10^−6^, in either European or African subsamples were removed. As a proxy for population stratification, we calculated principal components in PLINK using LD-pruned WGS data from GLOM and TUBE analyses separately.

### Single-SNP eQTL analysis

Single-SNP *cis*-EQTL (± 1Mb) analysis was performed with MatrixEQTL (Shabalin, 2012) adjusting for age, sex, batch, 4 genotype PCs, and PEER factors (Parts et al., 2011; Stegle et al., 2010). The optimal number of PEER factors was selected based on the maximum number of eGenes, as determined by TORUS (Wen, 2016), resulting in a variable number of PEER factors. Pearson correlation of SNP effect sizes for the top-ranked 5,000 genes were compared to other eQTL analyses including GLOM and TUBE from Gillies et al (Gillies et al., 2018) and Qiu et al. (Qiu et al., 2018) and GTEx kidney cortex (GTEx Consortium, 2017). To globally compare our GLOM and TUBE analyses to all GTEx tissues, we calculated principal components of *z*-scores from GLOM, TUBE and GTEx V8 eQTL analyses. For each gene, the SNP with the largest *z*-score across all studies was selected, resulting in one strong SNP association per gene.

### Enrichment of eSNPs in CREs of relevant kidney cell and tissue types

Enrichment of epigenetic annotations was estimated using TORUS (Wen, 2016) excluding the distance to TSS annotation. To compare the enrichment of eSNPs in CREs across different cell types, we controlled several potential confounding factors. First, we used peaks called from the same number of subsampled cells (N = 300). Second, we used the results of MatrixEQTL from the matched samples (N=219) between GLOM and TUBE. To analyze the baseline level of enrichment scores, we constructed two different matched control sets for the cell-type peaks: (1) peaks with the same number of target cell-type peaks randomly selected from the union of peaks covered in our kidney CRE map, and (2) randomly chosen genomic regions that have similar GC-contents and repeat fractions with the target cell-type peaks. To compare with different tissues, we applied our (optimized) pipeline to call the peaks from ENCODE DNase-seq datasets.

### Enrichment of eGene expression in relevant kidney cell and tissue types

We binarized snRNA-seq gene expression (GSE151302), where genes with non-zero expression in at least 2% of the cells were considered expressed for a given cell type. We performed a permutation analysis to compute cell-type specific gene enrichment of MatrixEQTL eGenes. To generate the null distribution, we randomized gene expression for each cell type, assuming a uniform distribution, and calculated the number of MatrixEQTL eGenes expressed for each cell type, repeating 1000 times. We calculated a *z*-score to quantify cell-type-specific gene enrichment in the snRNA-seq data set.

We conducted a similar permutation analysis using GTEx bulk RNA-seq data V8 obtained from the GTEx portal. To binarize gene expression, we ranked genes by median gene-level TPM for each tissue and classified genes ranked within the top 5000 as being expressed.

### SNP prior generation

Cell-type enrichment and biological relevance were considered when selecting cell-types to use for each tissue prior. For GLOM and TUBE separately, we generated base-pair resolution annotation files, where SNPs within ±300bp from the summit of ATAC-seq peaks from the union of selected cell-types were coded with a binary indicator. Using TORUS (Wen, 2016), we calculated enrichment of the CRE annotation along with distance to TSS, which was subsequently used to generate SNP priors for each gene.

### Multi-SNP eQTL analysis

We performed our integrative eQTL analysis with Deterministic Approximation of Posteriors (DAP) (Lee et al., 2018b; Wen et al., 2016) using genotype and expression data from NEPTUNE and TSS distances and our CRE-informed priors generated by TORUS. We adjusted for PEER factors (40 in GLOM, 50 in TUBE), age, sex, 4 genotype PCs, and RNA-seq batch. Gene-level Bayesian FDR methods were used to identify eGenes in each tissue and 95% credible sets were formed by summing ranked SNPs for each gene cluster.

### Comparison of fine-mapping results from different priors

We compared properties of the 95% credible sets to quantify fine-mapping resolution. For each credible set generated by each prior, we identified (1) the maximum snpPIP and (2) the number of SNPs in the credible set. Distributions from the uniform and integrative prior were compared with Wilcoxon rank sum tests.

### Computational prediction of regulatory effects

We generated deltaSVM scores to computationally predict the functional impact of SNPs (Lee et al., 2015). Open chromatin peaks of each cell type were used as a positive training set to build gkm-SVM models (Ghandi et al., 2014) as previously described, with some modifications. The LS-GKM (Lee, 2016) software with default parameter settings was used for training. To calculate comparable scores across cell-type models, (1) the top 100,000 peaks were used to train each model, and (2) deltaSVM scores were normalized per cell type using *z*-score based normalization of the distribution created by common SNPs with MAF > 1% in European ancestry from the 1000G project (Auton et al., 2015). *Z*-scores were transformed to probability scores for being functional variants using a logistic model trained by dsQTLs of lymphoblastoid cell line (LCL) (Degner et al., 2012) as a positive set and random SNPs with the control of GC contents and repeat fractions as a negative set. To aggregate the deltaSVM scores for GLOM and TUBE, we used the transformed scores of SNPs in peaks of the corresponding cell type and chose the best score per SNP among cell types whose CREs were used as the prior of corresponding eQTLs. We regarded the SNPs with aggregated score >0.5 as deltaSVM-positive in GLOM or TUBE compartments.

To test the significance of deltaSVM scores of the lead eSNPs, we identified random SNPs controlling for allelic frequency, distance from TSS, and the signal strength of open chromatin peaks (i.e., signal value of peak calling derived from MACS2). For per-SNP random control, we allowed 1%, 1000bp, and 1.0 as the residual error of the corresponding controlling variables, respectively.

### Colocalization

To test for colocalization of phenotype-associated SNPs and eSNPs from our tissue-specific eQTL analysis, we used fast enrichment estimation aided colocalization analysis, fastENLOC (Pividori et al., 2020; Wen et al., 2017), with default settings. The fine-mapped DAP results were converted to vcf format using the provided script summarize_dap2enloc.pl. *Z*-scores were extracted from trans-ethnic GWAS summary statistics (eGFR/UACR) (Teumer et al., 2019; Wuttke et al., 2019) and European 1000 Genomes project phase 3 version 5 samples (Auton et al., 2015) were used for the LD reference panel in all analyses. Of note, colocalization analysis with the European-only GWAS summary statistics yielded similar results.

When comparing our colocalization results to previous analyses, loci with RCP≥ 0.5 and matching tissue compartments were considered replicated, where GTEx cortex associations could match either GLOM or TUBE. To compare colocalization analysis with different priors, we used an RCP ≥ 0.2. We identified 1 SNP with the top SNP colocalization probability per colocalized eQTL cluster. When there were multiple top SNPs, we prioritized choosing SNPs that were in both the uniform and integrative prior and SNPs within ATAC peaks. To generate the expected number of SNPs within ATAC peaks, we randomly selected SNPs (100 x number of colocalized SNPs) controlling for the rank of the colocalized SNPs in the eQTL model. We calculated the mean overlap from 10 simulations for each tissue-trait pair. Enrichment was tested with a Binomial test of the observed overlap using the estimated expectations.

### Transcriptome-wide Association Analysis

Probabilistic transcriptome-wide association analysis (PTWAS) was used to test for causal relationships between GLOM and TUBE gene expression and complex kidney phenotypes – trans-ethnic meta-analyses of UACR (Teumer et al., 2019) and eGFR (Wuttke et al., 2019). Using the fine-mapped DAP results, glomerular and tubulointerstitial eQTL gene-SNP weights were calculated and formatted for GAMBIT gene-based testing using PTWAS helper scripts provided by the program authors (ptwas_builder, make_GAMBIT_DB.R). 1000 Genomes project phase 3 version 5 samples were used for the LD reference panel in all analyses. PTWAS_scan was run using default settings. Gene-level significance was adjusted to account for the multiple testing burden using two methods; q-value (Storey and Tibshirani, 2003) and a more conservative Bonferroni threshold. Genes with predicted pleiotropic effects or no strong instruments were excluded from our count of significant loci. We tested for pleiotropic effects and estimated effect sizes using the ptwas_est function.

### Gene validation in *Drosophila* nephrocytes

Genes for functional validation were selected based on the following criteria (1) causal association between gene and kidney phenotype in PTWAS analysis (FDR ≤ 0.05); (2) colocalization of eSNP associated with eGene and GWAS variants (RCP ≥ 0.5); (3) colocalization of eSNPs and relevant cell-type open chromatin; and (4) relative high expression levels of *Drosophila* homologs in the nephrocyte. A random subset of the qualifying genes was selected for functional follow up.

#### ANF-RFP uptake assay

Briefly, 10 virgin female flies from the MHC-ANF-RFP, Hand-GFP and Klf15-Gal4 transgenic lines were crossed to 5 male flies from UAS-RNAi transgenic lines of the targeted genes at 25°C. Pericardial nephrocytes of newly emerged adult flies (within 24 h of eclosion) were dissected and kept in artificial *Drosophila* hemolymph to assay RFP accumulation detected by fluorescence microscopy. For quantification of relative ANF-RFP fluorescence, 20 nephrocytes were analyzed from each of 3 flies per indicated genotype. T-tests were used to indicate significance differences from the control.

#### Dextran uptake assay

Flies carrying Hand-GFP and Klf15-Gal4 transgenes were crossed with flies carrying the UAS-RNAi transgenes at 25 °C. Dextran uptake was assessed in adult flies one-day post-emergence by dissection of pericardial nephrocyte in artificial *Drosophila* hemolymph and examination of the cells by fluorescence microscopy after a 20 min incubation with Texas Red labeled Dextran (10 kD, 0.02 mg/ml). For quantification of relative Dextran dye fluorescence, 20 nephrocytes were analyzed from each of 3 flies per indicated genotype. T-tests were used to indicate significance differences from the control.

### Luciferase reporter assay for allele-specific enhancer activity of rs80282103

Approximately 500-bp regions of DNA containing rs80282103 and rs11154336 were amplified from purified human genomic DNA (Promega, #G1521) by PCR using engineered restriction sites to allow directional cloning into the multiple cloning region of the pGL4.23[luc2/minP] luciferase reporter vector (Promega, #E841A). The resulting plasmids containing the insert in either “forward” or “reverse” orientation were confirmed by Sanger sequencing. Constructs containing the alternate alleles were obtained by performing Q5 site-directed mutagenesis (NEB, #E0554S). Primers used to amplify targets and perform site-directed mutagenesis are listed in **Table S9**. Each luciferase construct was co-transfected with pGL4.74[hRluc/TK] vector (Promega, #E692A), a *Renilla* luciferase control reporter, in HK-2 human proximal tubule cells at approximately 70% confluency in 96-well plates by using TransIT-2020 Reagent (Mirus, #5404), following the manufacturer’s protocol. Two separate transfections were performed with four technical replicates in each plate. Empty luciferase vector, pGL4.23[luc2/minP], was also transfected in quadruplicate as a control. Luciferase activity was quantified 48 hours after transfection using the Dual-Glo Reporter Assay System (Promega, #E2920) according to the manufacturer’s protocol. Luminescence signals were captured using a GloMax®-Multi+ Detection System (Promega) and normalized to *Renilla* luciferase readings for each well. We used linear regression with log transformed normalized luminescence adjusting for batch and orientation to test the allele effect on enhancer activity.

## Acknowledgements

NEPTUNE: The Nephrotic Syndrome Rare Disease Clinical Research Network III (NEPTUNE) is part of the Rare Diseases Clinical Research Network (RDCRN), which is funded by the National Institutes of Health (NIH) and led by the National Center for Advancing Translational Sciences (NCATS) through its Office of Rare Diseases Research (ORDR). NEPTUNE is funded under grant number U54DK083912 as a collaboration between NCATS and the National Institute of Diabetes and Digestive and Kidney Diseases (NIDDK). Additional funding and/or programmatic support for this project has also been provided by the University of Michigan, NephCure Kidney International and the Halpin Foundation. All RDCRN consortia are supported by the network’s Data Management and Coordinating Center (DMCC) (U2CTR002818). Funding support for the DMCC is provided by NCATS and the National Institute of Neurological Disorders and Stroke (NINDS).

DL is supported by Boston Children’s Hospital OFD/BTREC/CTREC Faculty Development Fellowship Award, NEPTUNE Career Development Fellowship, and a Manton Center Endowed Scholar Award.

A.C.O. is supported by the National Institutes of Health F32 Ruth L. Kirschstein Postdoctoral Individual National Research Service Award (DK122766).

The Genotype-Tissue Expression (GTEx) Project was supported by the Common Fund of the Office of the Director of the National Institutes of Health, and by NCI, NHGRI, NHLBI, NIDA, NIMH, and NINDS. The data used for the analyses described in this manuscript were obtained from the GTEx Portal on 03/16/2021.

This research is benefited by the data generated by the ENCODE consortium.

We thank Dr. Chakravarti for sharing unpublished adult kidney bulk ATAC-seq data.

The authors would like to acknowledge Boston Children’s Hospital’s High-Performance Computing Resources BCH HPC Clusters Enkefalos 2 (E2) and Massachusetts Green High-Performance Computing (MGHPCC) made available for conducting the research reported in this publication. Software used in the project was installed and configured by BioGrids (cite: eLife 2013;2:e01456, Collaboration gets the most out of software.)

---

| Tissue | Trait | Prior | Associations with RCP > 0.2 |  |  | % Overlap<br>(Expected)<br><sup>b</sup> | P-value <sup>c</sup> |
| --- | --- | --- | --- | --- | --- | --- | --- |
|  |  |  | Colocalized<br>SNPs <sup>a</sup> | SNPs in<br>ATAC<br>peaks | % Overlap<br>(Observed) |  |  |
| **GLOM** | eGFR | **Integrative** | 17 | 14 | 82.35% | 39.68% | 5.94x10<sup>-4</sup> |
| **GLOM** | UACR | **Integrative** | 32 | 22 | 68.75% | 42.65% | 3.69x10<sup>-3</sup> |
| **TUBE** | eGFR | **Integrative** | 69 | 56 | 81.16% | 62.34% | 1.08x10<sup>-3</sup> |
| **TUBE** | UACR | **Integrative** | 29 | 24 | 82.76% | 62.82% | 3.28 x10<sup>-2</sup> |
| **GLOM** | eGFR | **Uniform** | 4 | 1 | 25.00% | 8.45% | 0.30 |
| **GLOM** | UACR | **Uniform** | 13 | 0 | 0.00% | 7.71% | 0.62 |
| **TUBE** | eGFR | **Uniform** | 46 | 6 | 13.04% | 18.86% | 0.45 |
| **TUBE** | UACR | **Uniform** | 15 | 3 | 20.00% | 20.17% | 1.00 |

<sup>a</sup> One SNP per colocalized cluster, when > 1 SNP with same posterior probability, SNPs within ATAC peaks were prioritized
<sup>b</sup> Calculated from null distribution of SNPs matched for eQTL rank of colocalized SNPs (Methods)
<sup>c</sup> P-value from Binomial test of observed counts given expected % overlap

