## Supplemental Figures for "Mapping genomic regulation of kidney disease and traits through high-resolution and interpretable eQTLs"

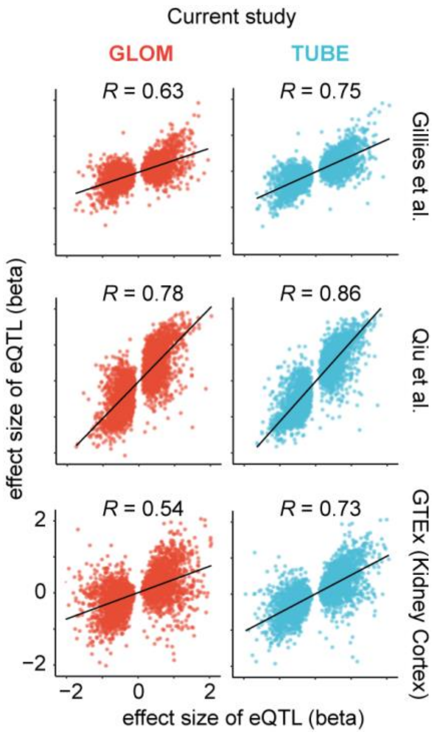


**Figure S1**. **Concordance of eQTLs across publicly available eQTL datasets.** Scatter plot and correlation of the effect sizes of GLOM and TUBE eQTLs paired with each kidney eQTL dataset.

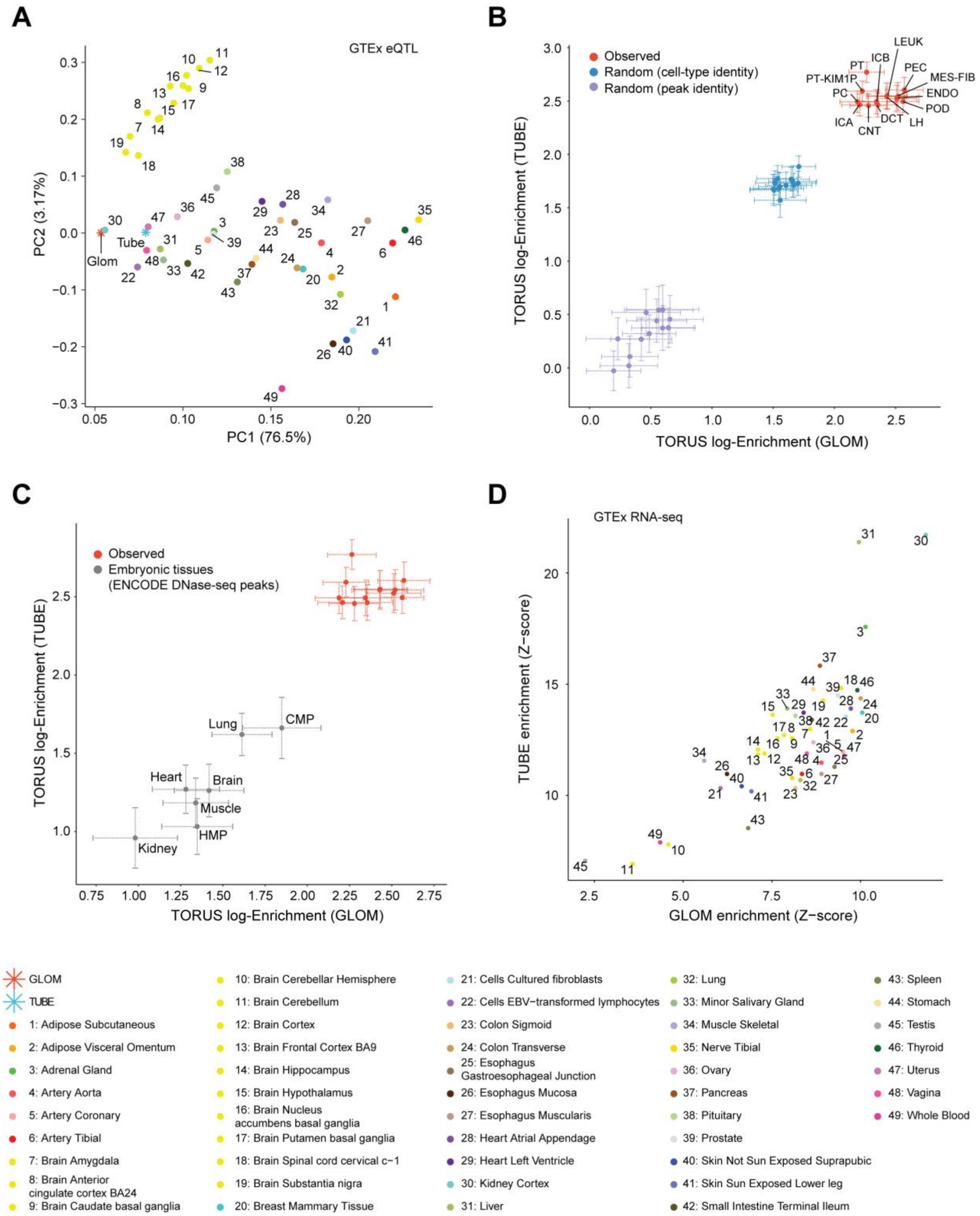


**Figure S2. The comparison of GLOM/TUBE eQTLs with eSNP/Genes and eQTL-association from GTEx/ENCODE samples. (A)** Principal component analysis of *z*-scores from independent eQTLs in GLOM, TUBE and GTEx tissues **(Methods). (B)** Red dots depict observed enrichment of our snATAC-seq peaks. Blue dots depict the enrichment of eSNPs in randomly sampled peaks from the union set of peaks across kidney cell types. Purple dots depict the enrichment of eSNPs in random regions of human genome (off peaks) that have the similar GC-contents and repeat fractions with the peaks of kidney cell types. **(C)** Enrichment analyses of eSNPs in open chromatin peaks of embryonic ENCODE tissues. **(D)** Enrichment analysis of eGenes in genes expressed in GTEx tissues.

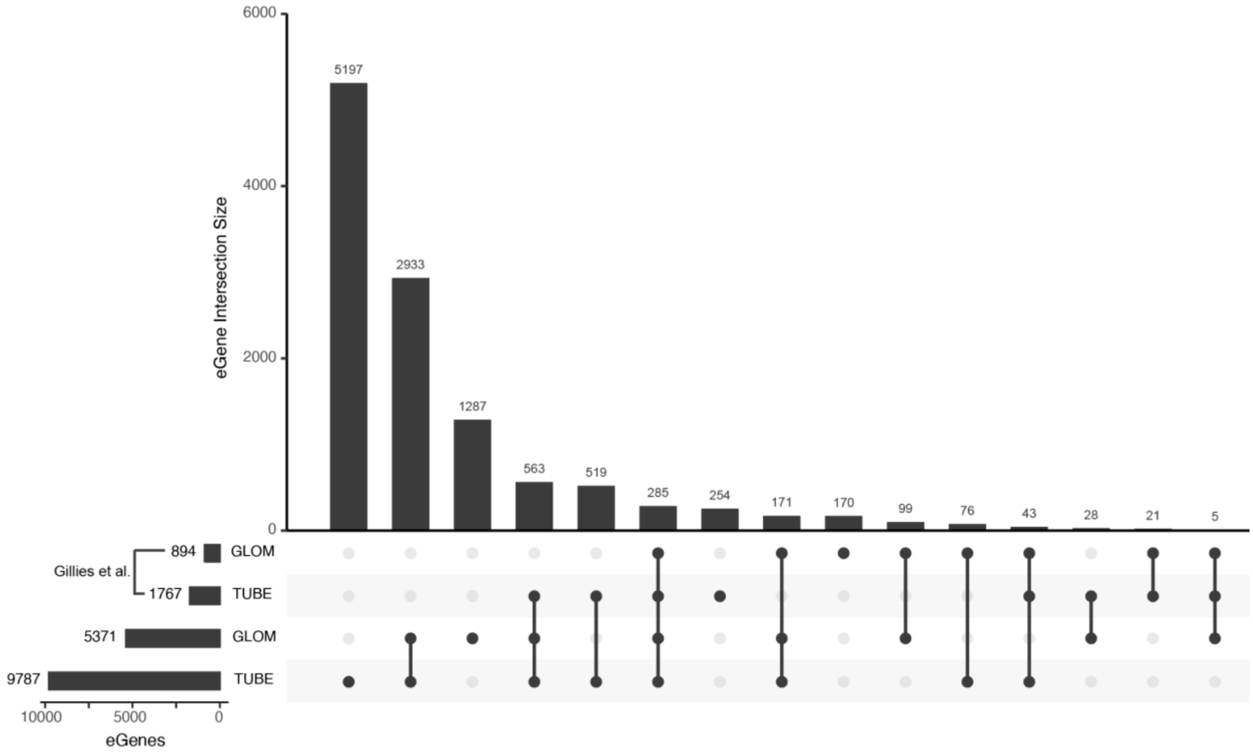


**Figure S3.** Upset plot comparing overlap of eGenes from array-based eQTL analyses (Gillies et al.) and CRE-informed GLOM and TUBE eQTL analyses.

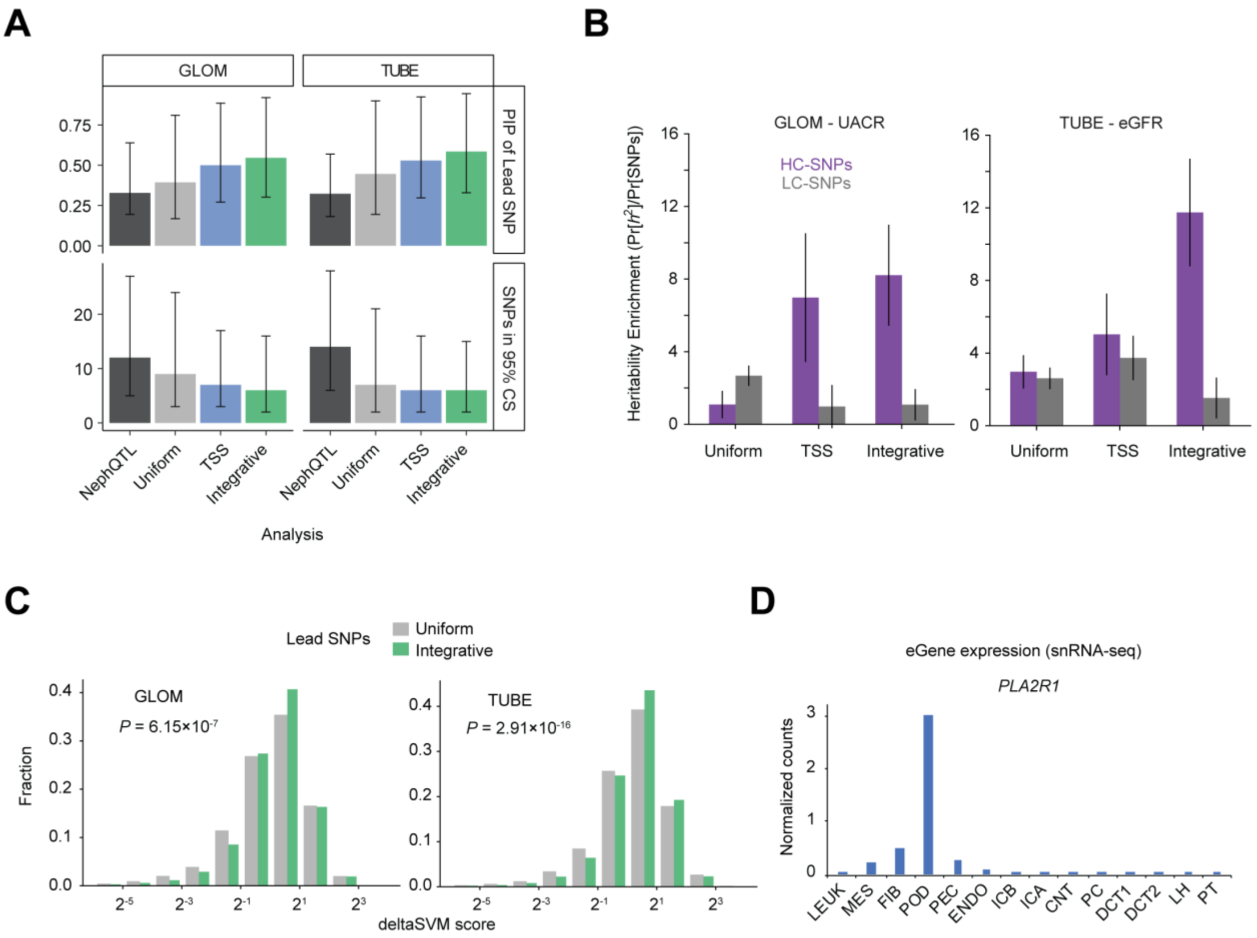


**Figure S4. Comprehensive analyses of fine-mapping quality across different priors. (A)** The distribution of the top SNPs' snpPIPs and number of SNPs forming the 95% credible set. Compared to **Figure 4A**, NephQTL (Gillies et al., 2018; TSS-only prior) and current eQTLs with TSS-only informed prior are added. **(B)** S-LDSC enrichment analysis of GLOM-UACR and TUBE-eGFR eQTL-trait combinations, GLOM/TUBE eQTLs with the three different priors as shown in **Figure 4B**. **(C)** Comparisons of deltaSVM scores between lead SNPs from uniform and integrative priors as shown in **Figure 4C**. **(D)** Gene expression of *PLA2R1* in snRNA-seq data normalized by genes and cell counts.

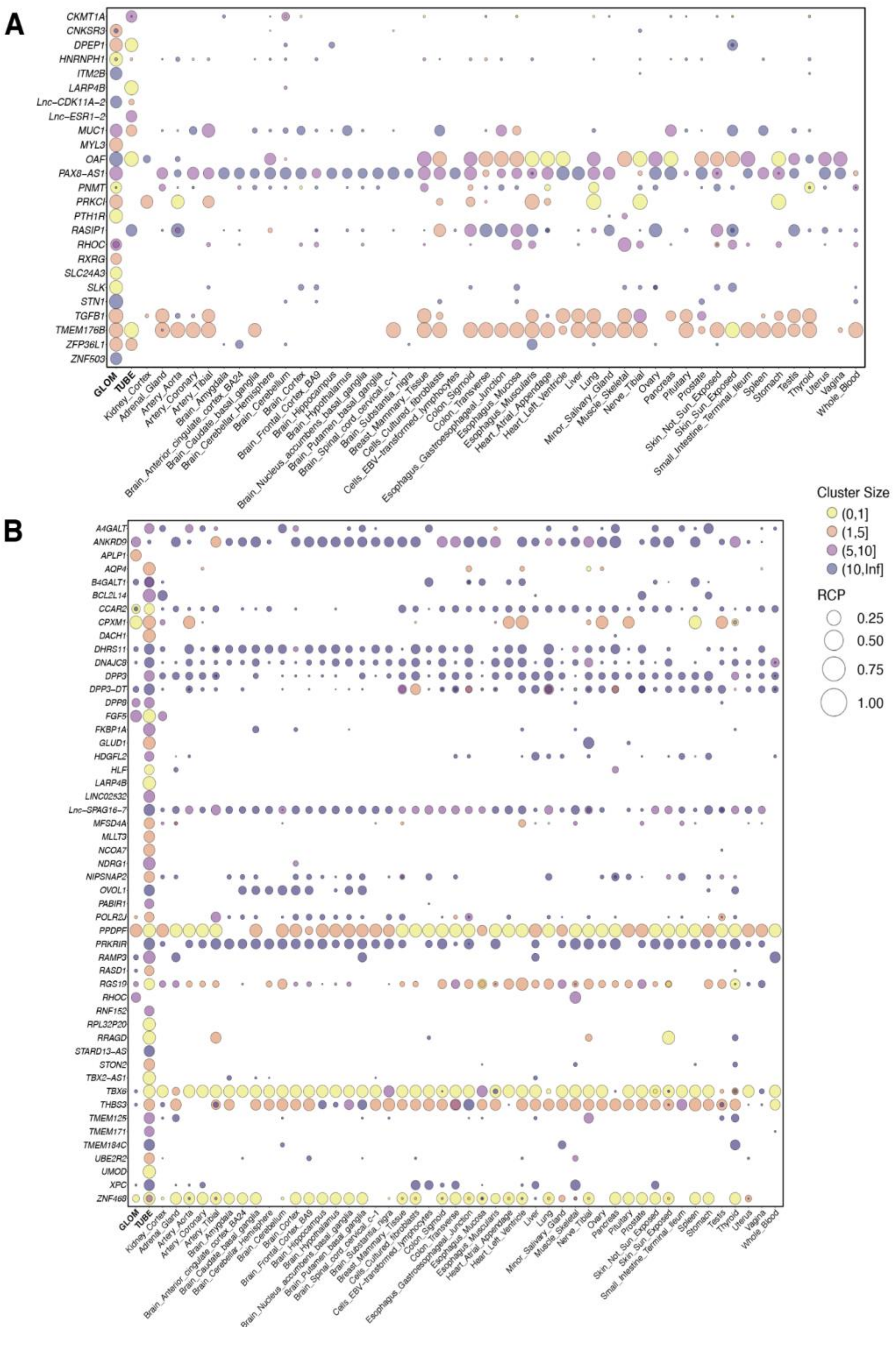


**Figure S5. Colocalized genes using integrative prior.** Each bubble plot shows GLOM and/or TUBE eGenes colocalized with **(A)** UACR and **(B)** eGFR GWAS loci (≥1 cluster with Regional colocalization probability (RCP) ≥ 0.5) along with GTEx results colocalization results. Each circle represents an eQTL cluster, and its diameter and color depict the RCP scores and the cluster size (number of eSNPs), respectively.

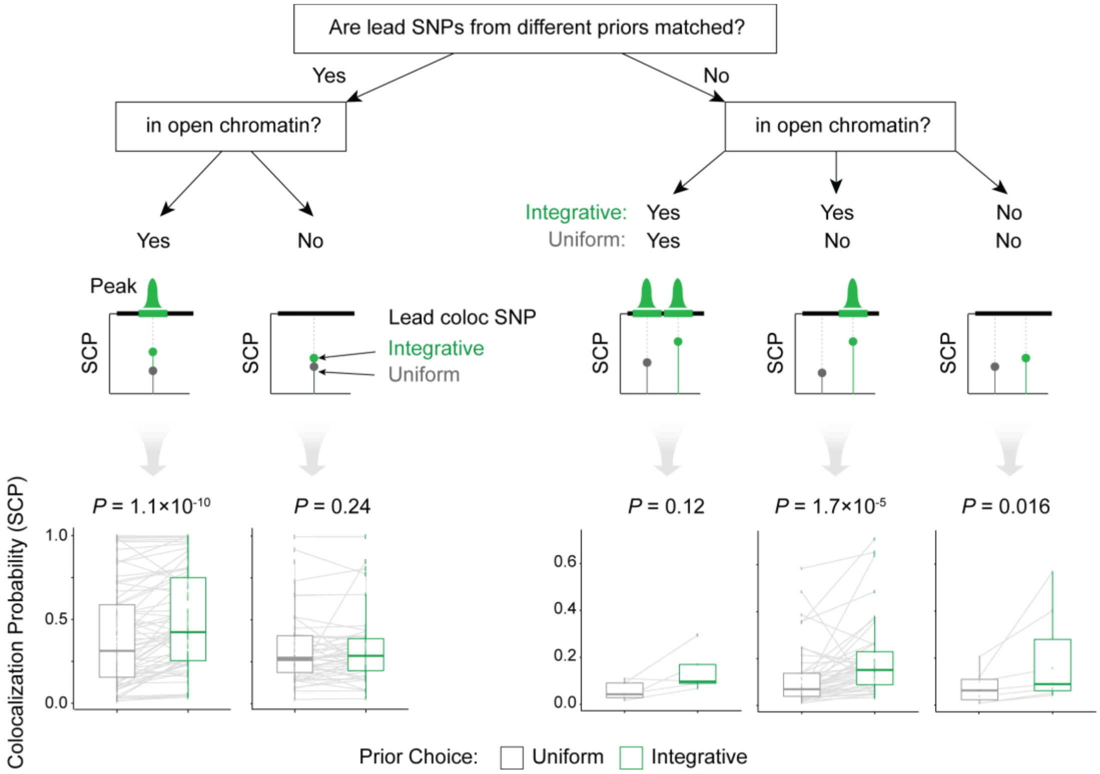


**Figure S6.** SNP colocalization probability of lead SNP (highest probability) for each colocalized cluster. Association tests are stratified by open chromatin status and the prior used to discover the gene-SNP pair. All four analyses (GLOM-eGFR, GLOM-UACR, TUBE-eGFR, TUBE-UACR) are combined for comparisons. To compare the increase/decrease in colocalization probability with respect to the prior choice, lead SNPs where the colocalized eSNPs share the same eGene are paired. To test the statistical significance paired-sample Wilcoxon test was used. Notably, there are no SNP pairs in which the lead SNP from integrative prior is in closed chromatin and the lead SNP from uniform prior is in open chromatin.

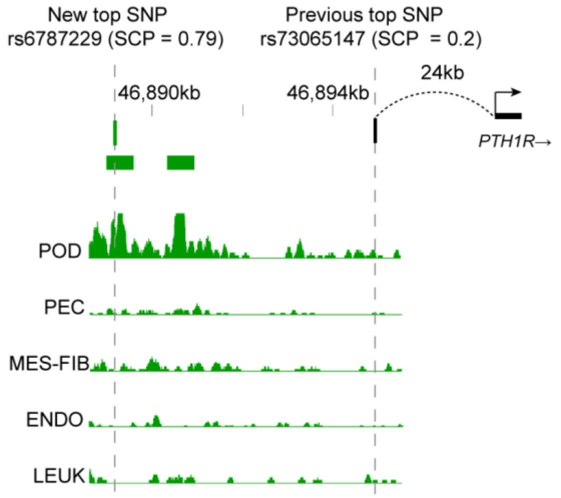


**Figure S7.** Refined top colocalized SNP for GLOM *PTH1R* and UACR. Black SNP is from Wuttke et al. colocalization analysis using array-based GLOM eQTLs (Gillies et at.). Vertical dashed lines connect the lead SNPs to their genomic coordinate with an open-chromatin perspective. Horizontal and vertical bar plots depict the genomic range of open chromatin peaks and the pile-up of snATAC-seq reads on the relevant cell types.

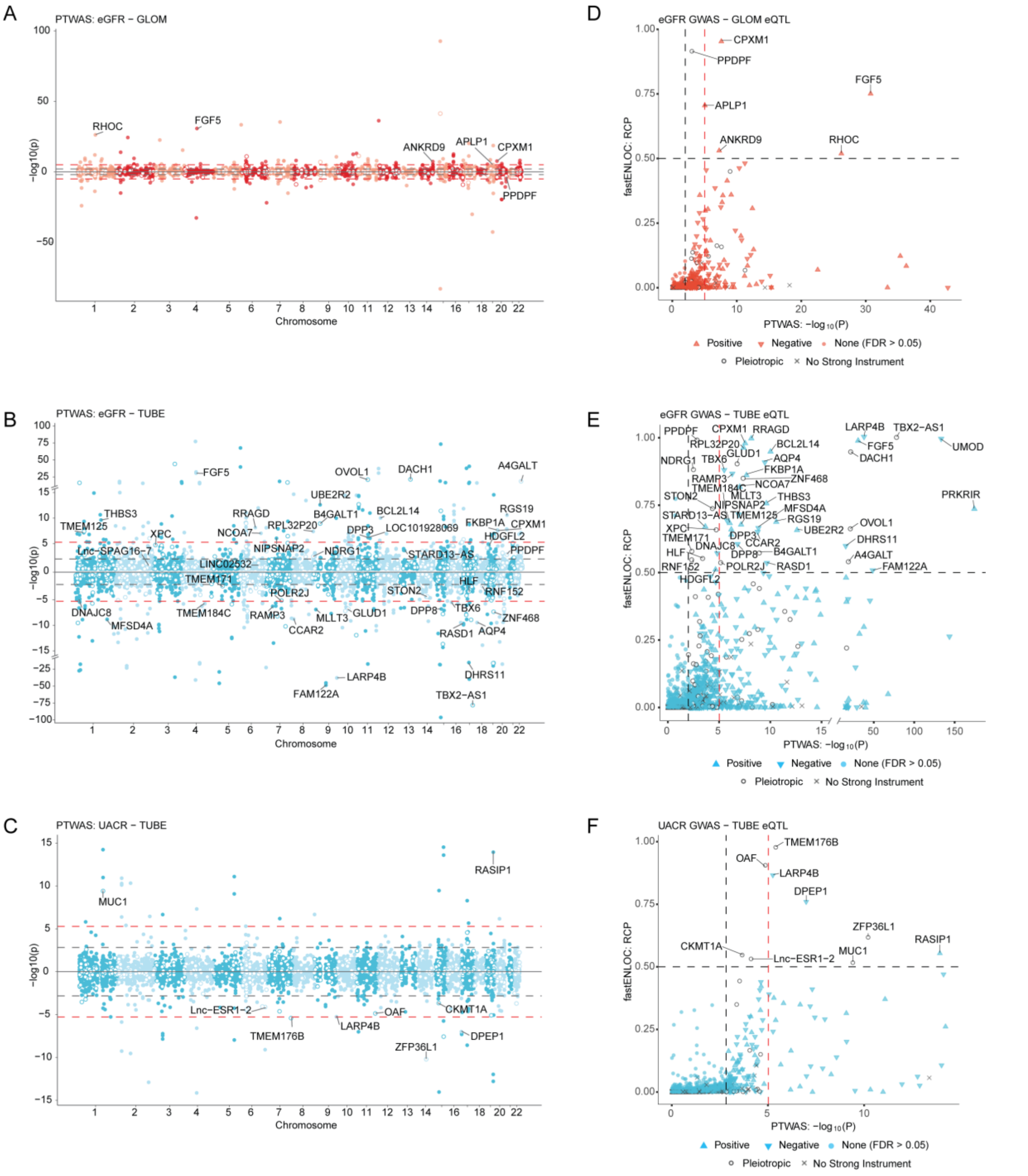


**Figure S8.** PTWAS Miami plots for **(A)** eGFR - GLOM, **(B)** eGFR - TUBE, and **(C)** UACR - TUBE. Each dot represents a gene, and the genes potentially confounded by pleiotropic effects are indicated with an open circle. Genes with no strong instrument are excluded. Dashed lines indicate thresholds using two multiple testing correction methods, *q*-value ($q \leq 0.05$; light gray) and Bonferroni ($P \leq 9.52\times10^{-6}$; red). Genes with regional colocalization probability (RCP) ≥ 0.5 from the corresponding colocalization analysis are labeled. **(D-F)** Scatter plots of RCP of top colocalized clusters from fastENLOC and corresponding PTWAS associations for each eGene. The shape of points depicts the type of effect of the PTWAS association, as shown in the figure. Genes with no strong instrument are excluded.

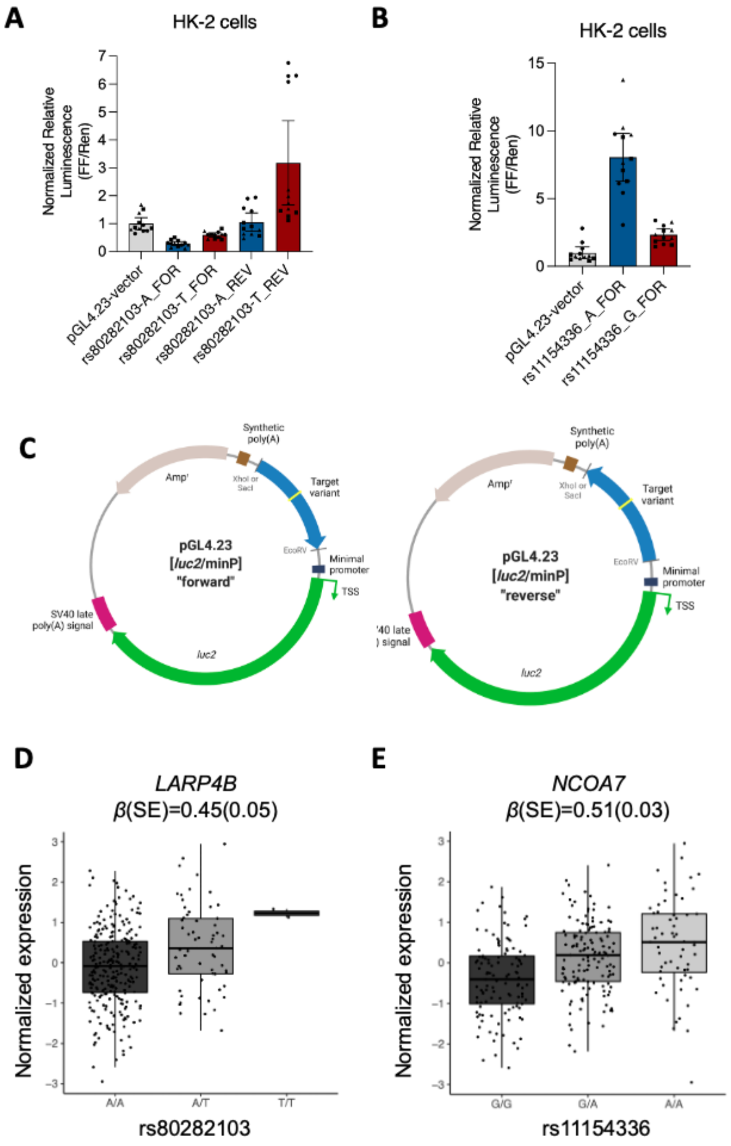


**Figure S9. (A-B)** Reporter assay in HK-2 cells testing the allele-specific enhancer activity (A) rs80282103 and (B) rs11154336. Results are given as ratios of firefly to *Renilla* luciferase activity. Results shown from three independent experiments with quadruplicate measurements in each repeat. *P*-value from linear regression model with log transformed relative luminescence = rs80282103: $5.99\times10^{-9}$, rs11154336: $1.05\times10^{-8}$. Note that only forward orientation is available for rs11154336. (C) Diagrams for the luciferase constructs. (**D-E**) TUBE eQTL boxplots of (D) *LARP4B*-rs80282103 and (E) *NCOA7*-rs11154336.
